# Genetic basis of partner choice

**DOI:** 10.1101/2025.02.03.636375

**Authors:** Qinwen Zheng, Sjoerd van Alten, Torkild Hovde Lyngstad, Edoardo Ciscato, Zhongxuan Sun, Jiacheng Miao, Yuchang Wu, Stephen Dorn, Boyan Zheng, Alexandra Havdahl, Elizabeth C. Corfield, Michel Nivard, Titus J. Galama, Patrick Turley, Pierre-André Chiappori, Jason M. Fletcher, Qiongshi Lu

## Abstract

Previous genetic studies of human assortative mating have primarily focused on searching for its genomic footprint but have revealed limited insights into its biological and social mechanisms. Combining insights from the economics of the marriage market with advanced tools in statistical genetics, we perform the first genome-wide association study (GWAS) on a latent index for partner choice. Using 206,617 individuals from four global cohorts, we uncover phenotypic characteristics and social processes underlying assortative mating. We identify a broadly robust genetic component of the partner choice index between sexes and several countries and identify its genetic correlates. We also provide solutions to reduce assortative mating-driven biases in genetic studies of complex traits by conditioning GWAS summary statistics on the genetic associations with the latent partner choice index.

## Introduction

Partners resemble one another in various characteristics, including sociocultural, physical, socioeconomic, health, and personality traits^1–6^. Partner selection based on resemblance is often referred to as assortative mating in a broad literature. The dynamics of partner choice and assortative mating have received considerable attention across both social and biological sciences^7^. For example, assortative mating may contribute to explaining the recent reversal of the gender gap in higher education^8^, the increase in women’s labor supply^9^, the rise in socioeconomic inequality in developed countries both within and across generations^10–15^, and the comorbidity of various psychiatric traits^4,16^. In genetics, positive assortative mating decreases phenotypic variance in the population with each consecutive generation^17^ and may induce long-range linkage disequilibrium (LD), a correlation in otherwise independent genetic variants^5^. It is known that spouses exhibit similarity for genetic factors associated with various phenotypes, such as educational attainment (EA), height, and body-mass index (BMI)^5,16,18–21^. Using independent individuals in biobank cohorts, several studies have identified long-range LD among genetic variants associated with these phenotypes^22–25^. As a result, assortative mating may induce a genetic correlation between phenotypes that do not share the same causal genetic pathways^26^. Ignoring assortative mating can thus lead to serious biases in estimates of variant effects on complex traits, heritability, and genetic correlation^26–31^. However, few studies have explored approaches to remedy these issues to date^32^. Indeed, most genetic studies of complex human traits assume random mating.

Previous genomic studies on human assortative mating have two major limitations. First, most existing studies quantified genetic similarity (between family members or between independent chromosomes within the same individual) at the polygenic score (PGS) level^22,23,25^. These scores effectively quantify the cumulative effect of thousands to millions of genetic variants on a trait of interest, but the use of PGS in assortative mating research is problematic for at least two reasons: (a) PGS are typically measured with considerable error so that spousal correlations will likely be biased and (b) individual PGS are created to predict a single phenotype, and thus investigators must choose specific phenotypes (e.g. height) rather than broader assortative patterns. Methodology for directly identifying the possibly unmeasured traits driving assortative mating has been lacking. Variant-based approaches to assessing assortative mating have also been problematic. Since investigators do not yet understand how individual genetic variants exert pressure on assortative mating, existing efforts use hypothesis free scans across the genome to estimate spousal correlation at each single variant, which is underpowered^33^. Further, the lack of a remedy to control for assortative mating-driven biases may have contributed to the very limited consideration of assortative mating in human genetic studies. This paper proposes a solution to this problem by leveraging interdisciplinary methodological advances from statistical genetics and economics and by conducting sufficiently powered analyses across multiple global cohorts.

We adopt a partner matching model from economics and show that associations between the genome and a partner choice index can be uncovered by performing genome-wide association studies (GWAS) on various spousal traits. Subsequently, we perform such GWAS on spousal EA, income, height, and BMI using more than 100,000 opposite-sex couples of European ancestry from four large cohorts containing genome-wide genetic data: UK Biobank (UKB), the Health and Retirement Study (HRS) from the US, Norwegian Mother, Father and Child Cohort Study (MoBa), and the Lifelines Cohort Study (Lifelines) from the Netherlands. BMI, EA, and income have previously been used to inform the partner choice index in the economics literature^34^. We also include height in the analysis due to its evident involvement in assortative mating^5,22,35,36^. We extract the shared genetic component across multiple spousal traits to represent the genetics of the partner choice index and demonstrate a broadly robust genetic component across countries, environments, and sexes. Our results obtained using genetic data provide empirical evidence for the index factor model^37^ used in economics and shed important light on the genetic underpinnings of partner choice. Further, we show that our GWAS results for the partner choice index can be used to correct for assortative mating-driven inflation in complex trait genetic applications.

## Results

### Model setting and key intuitions

Following a broad literature in economics, we adopt a matching model in which we assume that the (multi-dimensional) individual preferences in partner choice can be summarized by a one-dimensional index, which is a function of many characteristics^37^. We use a linear-model setting as a working assumption and discuss how to relax this assumption in the **Supplementary Note.**

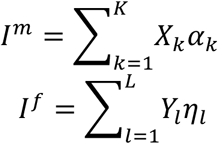

Here, *I^m^* and *I^f^* are the indices for males and females, respectively. *X*_1_, *X*_2_, …, *X_K_* (*Y*_1_, *Y*_2_, …, *Y_K_*) are the set of male (female) characteristics that determine the index; *K* (*L*) is the total number of male (female) characteristics. This model assumes that an individual’s characteristics matter for partner preference only through the index (**Figure 1**). Intuitively, this is suggesting that if two men (women) have the same values of *I^m^* (*I^f^*), they are equivalent in terms of their desirability in the context of partner selection, even though *I^m^*(*I^f^*) can result from different individual characteristics. In the economics literature, this index has been referred to as “attractiveness in the marriage market”. But we note that this index is different from physical attractiveness and is known to involve many social, cultural, and economic characteristics^34^. For example, the index may also operate through individuals self-selecting into environments that are conducive to finding a suitable spouse. Additionally, with respect to the terminology used to describe the process of partner selection, we aim to balance accurately reflecting contemporary social realities with facilitating clear connections to established literature, particularly within the field of economics where “marriage market” is a prevalent term in matching theory and analysis. The term and the associated analysis is an extension of labor market analysis and household decision making models in economics^38,39^, for example, where firms and workers search for and enter into employment matches based on their characteristics. Likewise, a marriage market is intended to represent the process by which individuals find partners and mutually agree to form a stable relationship—defined as a relationship that is either co-residential, involves the sharing of at least one common child, or both. Further, although real-world marriage markets are much more complex, empirical evidence in the economics literature shows that the low-dimensional index model is a reasonable approximation consistent with real-world data^37,40,41^. We discuss this further below and in the **Supplementary Note**.

**Figure 1.**
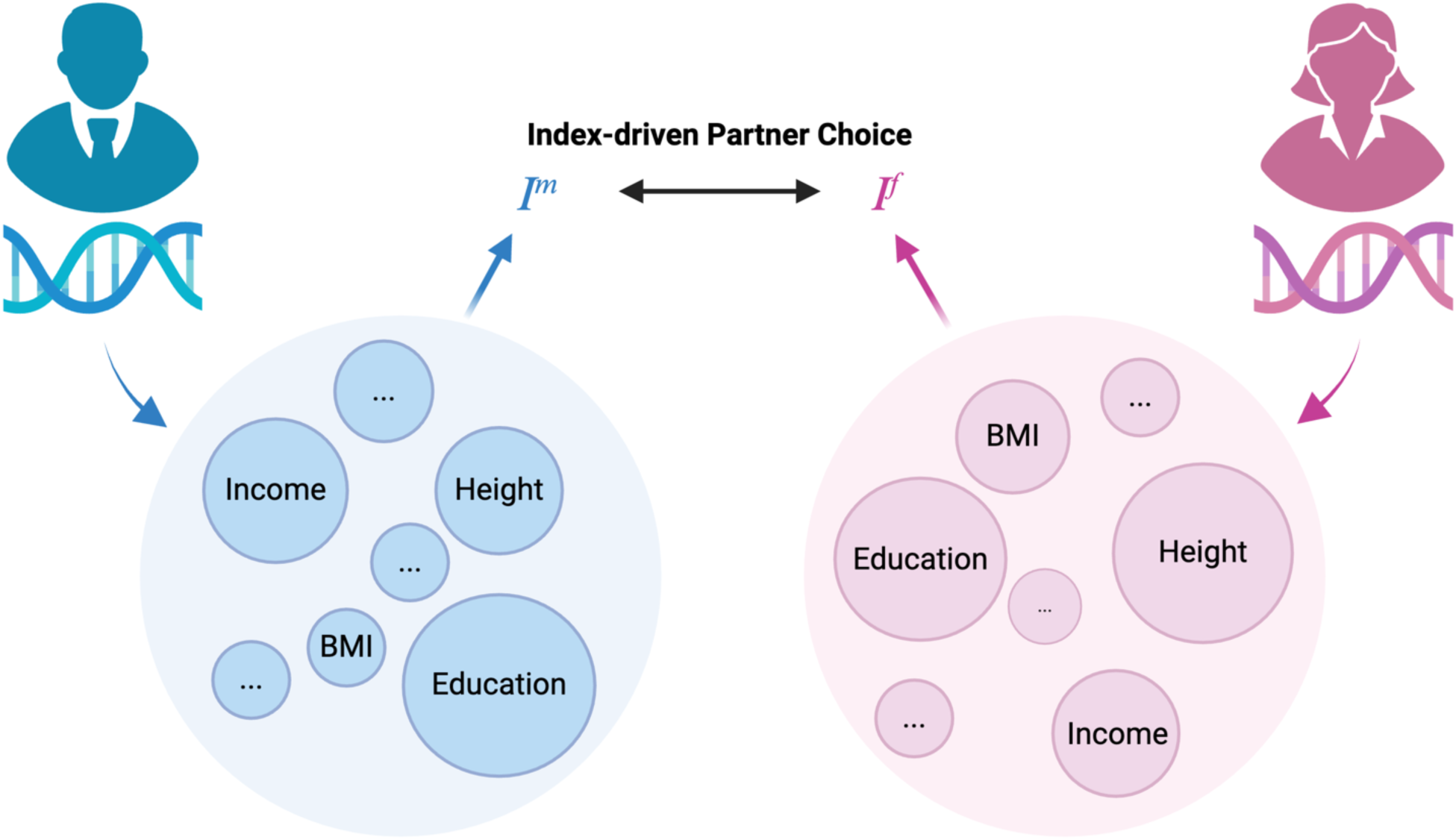
An Illustration of index-driven partner choice. Indices *I^m^* and *I^f^* respectively represent a man’s and a woman’s “attractiveness” in the marriage market. Height, income, BMI, and education are determinants of *I^m^* and *I^f^* that are observed in our study. Albeit different with respect to such traits, two individuals are equally attractive if they share the same index value. Further, our model allows for an unknown number of unobserved traits (here denoted as “…”) to affect *I^f^* and *I^m^*. BMI: body-mass index. This figure is created via BioRender.

Our primary goal in this study about the genetic basis of partner choice, is to perform a GWAS on this index. However, a main challenge is that *I^m^* and *I^f^* are latent, making it impossible to directly perform a GWAS.

Following convention in complex trait genetic studies, we assume that the individual characteristics *X_k_* and *Y_l_* are affected by both genetics and environments.

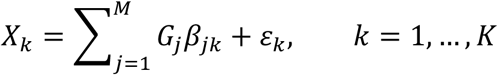

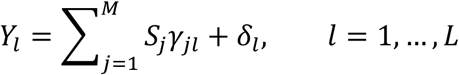

Here, *G_j_* (*S_j_*) is the j-th single-nucleotide polymorphism (SNP) of a male (female) individual; *β_jk_* (*γ_jl_*) is its effect size on the k-th male (l-th female) characteristic; *ε_k_* and *δ_l_* denote the non-genetic components underlying these characteristics; *M* is the total number of SNPs. We note that *G* = (*G*_1_, …, *G_M_*) and *S* = (*S*_1_, …, *S_M_*) are the same set of SNPs. We use different notations for male and female genetic variables just for convenience and clarity in later derivations.

Our main results depend on two key assumptions which are discussed in greater details in the **Supplementary Note**:

- ***Separability*** holds if there exists a metric *θ* (*τ*) that entirely subsumes the role of genetics in the male (female) index. In other words, if two men share similar non-genetic components *ε* = (*ε*_1_, …, *ε_K_*) but have different SNP genotypes *G* and *G*′, they are equally attractive, i.e. they have the same index value *I^m^*, as long as *θ*(*G*) = *θ*(*G*^’^). Similarly, two women with similar non-genetic components *δ* = (*δ*_1_, …, *δ_L_*) but different SNPs *S* and *S*’ are equally attractive as long as τ(*S*) = τ(*S*’). We refer to *I^m^* = *θ*(*G*) and *I^f^*= *τ*(*S*) as the genetic components of the main indices *I^m^* and *I^f^*.
- the linear model setting, it is easy to see that the separability assumption holds. That is, genetics only matter for partner choice through the genetic components:

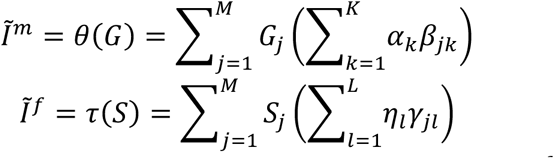

We illustrate below that there is no need to distinguish *I^m^*(*I^f^*) and *II^m^* (*I^f^*) for the purpose of genetic association analysis under linear models.

- ***Conditional Independence*** holds if the distribution of *ε* (*δ*) is independent of *G* (*S*) conditional on *I^m^*(*I^f^*). This means that the conditional distribution of non-genetic traits *ε* is identical for any two men with SNPs *G* and *G*′as long as *θ*(*G*) = *θ*(*G*′). The same applies to women. We note that, in many complex trait genetic studies^42^, it is standard to assume independence of *G* (*S*) and *ε* (*δ*). Here, we only require a weaker assumption of conditional independence.

A key property of this framework which makes empirical analyses possible in this study is that, under the separability and conditional independence assumptions, we have:

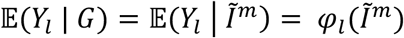

That is, the conditional expectation of the l-th characteristic of the partner given an individual’s genetics *G* is a function of the individual’s index *I^m^*. A similar conclusion holds true for the female index *I^f^*. A key implication of this is that, although it is challenging to directly measure *I^m^*(*I^f^*), its genetic components *I^m^*(*I^f^*) can be identified (up to an increasing transformation) using couples’ observable characteristics. That is, a characteristic *Y_l_* of the wife (e.g., EA) is informative on the husband’s value of the latent partner choice index, and vice versa. Even more importantly, the relative contribution of genetic variant *G_j_* on the index *I^m^* (i.e., *∂I^m^*/*∂G_j_*) can be empirically studied.

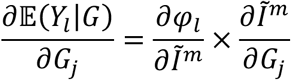

That is, *∂Ĩ^m^*/*∂G_j_* can be obtained from empirically observable *∂*𝔼(*Y_l_*|*G*)/*∂G_j_* up to a scalar *∂φ_j_*/*∂Ĩ^m^* which does not depend on *j*. Since *∂I^m^*/*∂G_j_* = *∂Ĩ^m^*/*∂G_j_* in the linear model setting, this means that genetic contributions to the index *I^m^* can be empirically estimated. In the context of genetic association analysis, if we further assume *∂φ_j_*/*∂Ĩ^m^* is constant across individuals, we can regress wife’s trait *Y_l_* (e.g., EA) on husband’s SNP *G_j_*, and this will give us the SNP association with the husband’s index *I^m^*, up to a scalar that does not depend on the choice of SNP. Therefore, a GWAS on wife’s EA is (almost) equivalent to our intended GWAS on *I^m^* and identifies the genetic component of *I^m^*. The choice of the wife’s trait is arbitrary (as long as it contributes to the index) and the GWAS is robust to this choice, which provides a practical advantage in that we do not need to observe all, or even many, traits of the wife. However, if we do observe multiple characteristics of the wife, we can perform multiple GWAS on these characteristics (e.g., income, height, BMI). These GWAS, performed on different spousal traits, are each, essentially, a GWAS of the partner choice index up to a scaling factor difference. In practice, we employ factor GWAS techniques^43^ to identify the shared genetic component across multiple spousal traits and use it to represent the GWAS of the partner choice index. In the **Supplementary Note**, we show that this strategy can identify genetic associations with the index even if the single index assumption is violated by trait-specific matching (i.e., the index and individual characteristics jointly determine partner choice; **Supplementary Figure 1**).

### GWAS on spousal outcomes

We performed GWAS on four spousal traits: height, BMI, EA, and income (sample size ranging 151,504-203,617) using individuals of European ancestry from UKB, HRS, MoBa, and Lifelines (**Supplementary Table 1** and **Supplementary Figures 2-5**). In the initial analysis, we ignore sex differences and maximize our sample size to identify genetic associations with a partner choice index *I* that is shared between males and females. GWAS were performed in each cohort separately, adjusting for sex, age, 20 principal components, and cohort-specific batch covariates. Then, a fixed-effect meta-analysis was conducted. We identified 1, 3, and 10 genome-wide significant loci for spousal height, BMI, and EA, respectively (**Supplementary Table 2** and **Supplementary Figures 6-8**). Within-sibling GWAS produces estimates of direct genetic effects that are robust to various types of confounding^44^, but is not sufficiently powered in this study to identify SNP-level associations (**Methods**). However, 10 of these 14 loci showed consistent effect directions in within-sibling analysis despite the small sample size (**Supplementary Tables 2-3**).

All four traits had SNP-based heritability estimates that are significantly greater than 0 (**Figure 2A** and **Supplementary Table 4**), with spousal EA showing the highest heritability (h^2^ = 0.08, se = 0.004). These estimates are similar to previous reports based on UKB alone^45^. As a comparison, we also performed GWAS on own outcomes for the same set of traits in the same study sample (**Supplementary Table 1**). Heritability of spousal EA was about half of the estimate for own EA (h^2^ = 0.17, se = 0.006). For height, BMI, and income, own traits showed substantially higher heritability compared to spousal traits (**Figure 2A**). Based on the partner choice index model, we expected high genetic correlations across spousal traits. Indeed, we found substantially higher genetic correlations among spousal traits compared to own traits, showing an average 2.57-fold increase across trait pairs (**Figure 2B** and **Supplementary Table 5**). For example, spousal BMI and income had a genetic correlation of -0.92 (se = 0.14), while the genetic correlation of own BMI and income was -0.27 (se = 0.03). The intuition of this is that, while BMI and income are very different characteristics, they are both (similarly) informative on the partner’s index in the marriage market. This pattern was consistently observed in all trait pairs, which provides empirical support for the index model.

**Figure 2.**
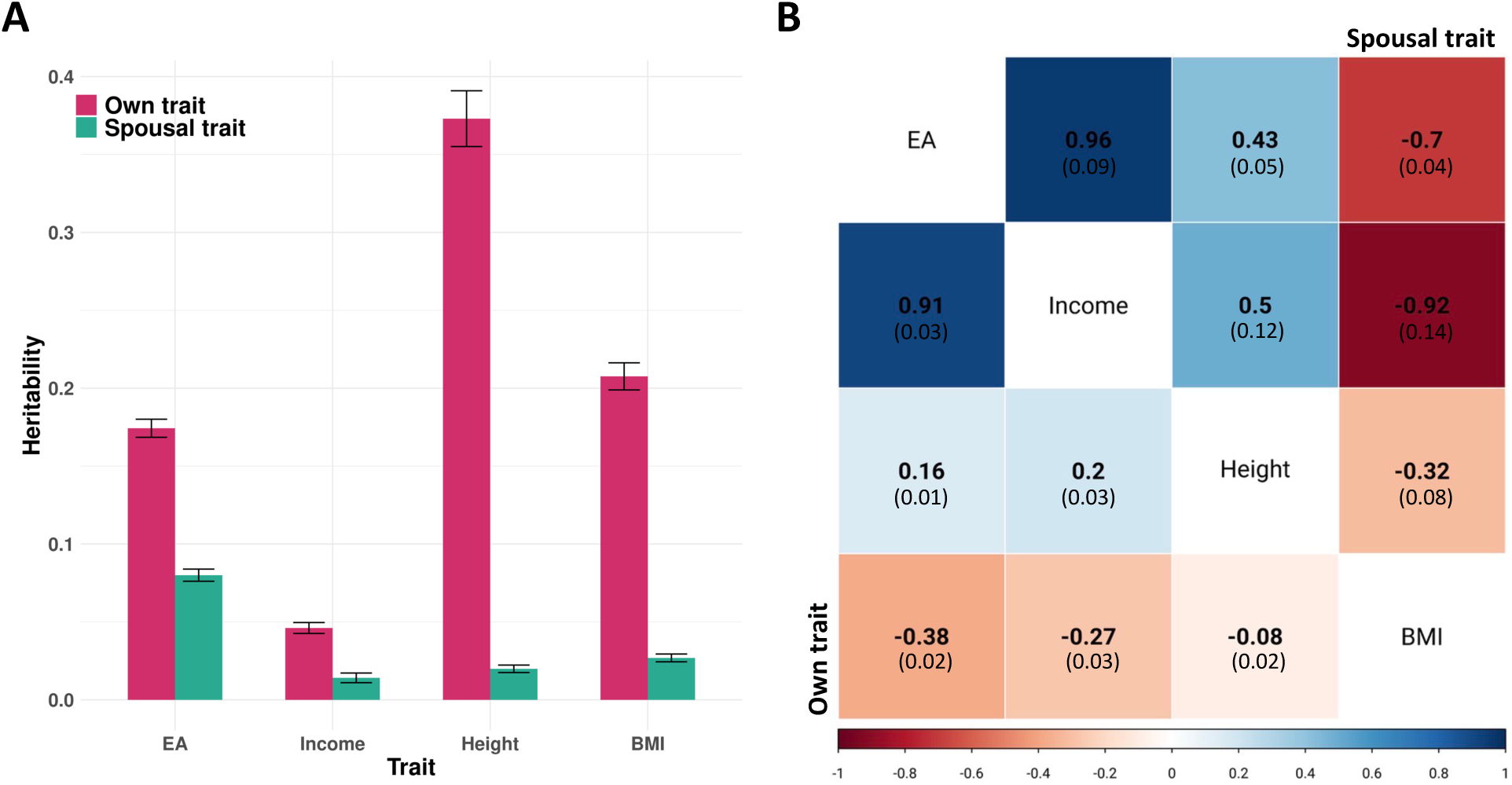
Heritability and genetic correlation estimates based on GWAS of own and spousal outcomes. **(A)** Heritability estimates obtained from GWAS of spousal and own traits. Intervals indicate standard errors. **(B)** Pairwise genetic correlations for four complex traits. Standard errors are shown in the parentheses. The upper triangle shows the genetic correlations between spousal traits and the lower triangle shows that of own traits.

### Genetic associations with the partner choice index

Although the genetic correlations among spousal traits are high, they are not 1 or -1 as the single index model implies. We employed a genomic structural equation model (GenomicSEM^43^) with a single factor to identify the shared genetic component underlying spousal traits (**Figure 3A**; Comparative Fit Index = 1 and Standardized Root Mean Square Residual = 0.021, indicating good model fit). As we demonstrate in the **Supplementary Note**, this approach identifies genetic associations with the index even when trait-specific matching also influences partner choice in addition to the index. We performed factor GWAS using summary statistics of four spousal traits as the input (**Methods**). We refer to this GWAS as the index GWAS throughout the paper.

**Figure 3.**
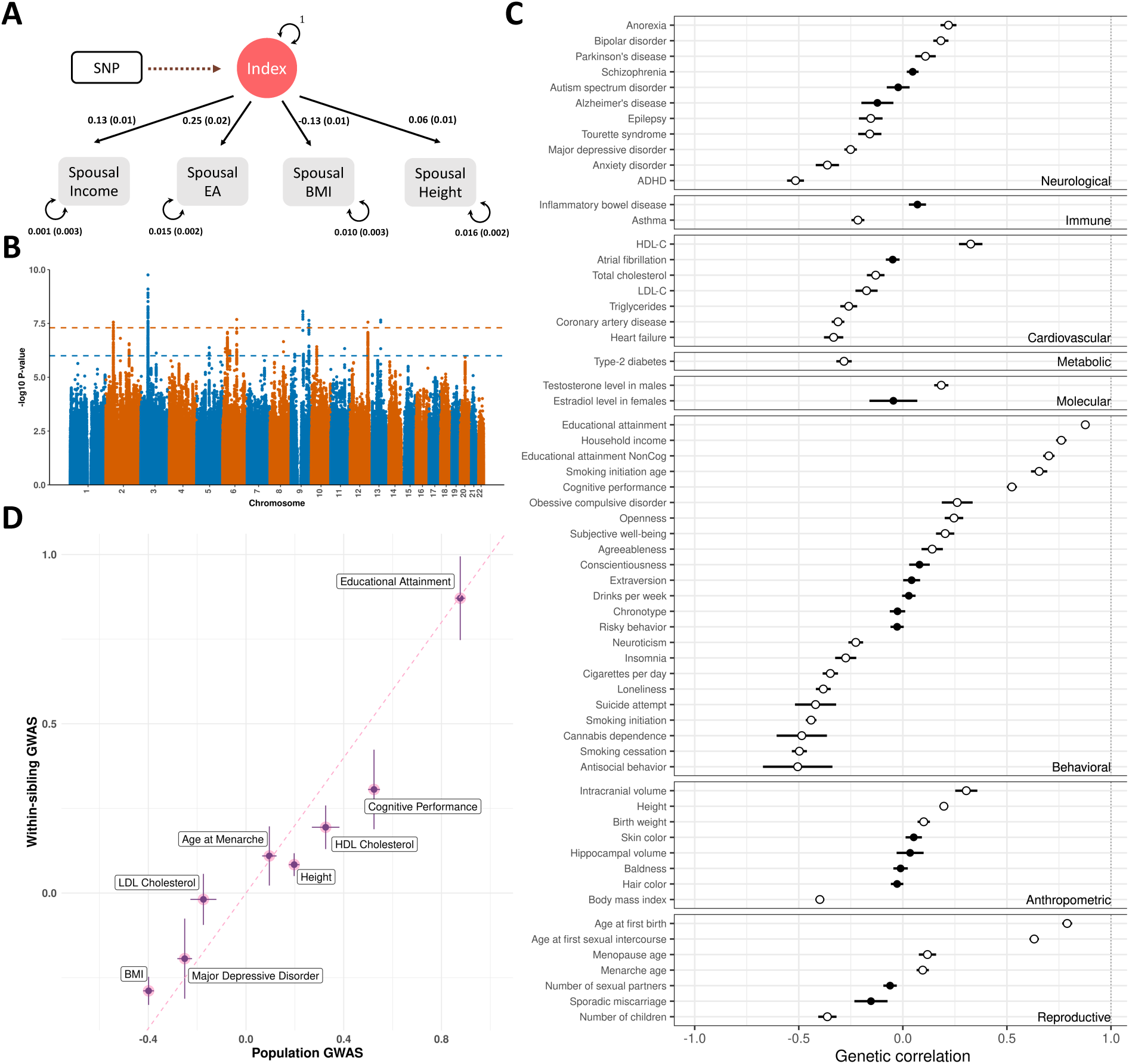
GWAS results for the partner choice index. **(A)** Single-factor GenomicSEM model for four spousal traits. Factor loadings on each individual spousal trait were estimated without including SNPs in the model. Standard errors are shown in the parentheses **(B)** Manhattan plot of the index GWAS. The orange horizontal line indicates genome-wide significance (p < 5e-8), the blue line indicates a suggestive significance level at 1e-6. **(C)** Genetic correlations between the index GWAS and 64 complex traits. Intervals indicate standard errors. Hollow circles indicate significant correlations at false discovery rate (FDR) < 0.05. **(D)** Genetic correlations of the partner choice index with population GWAS (X-axis) and within sibling GWAS (Y-axis) of eight complex traits. Index genetic associations were obtained from population-based analysis. In (C) and (D), each interval indicates ±1 standard error.

We identified seven genome-wide significant loci associated with the partner choice index (**Figure 3B**, **Supplementary Table 6**, and **Supplementary** Figure 9). The most significant SNP (i.e., rs7613360-C; p = 1.7e-10) is a known correlate of higher cognitive ability^46,47^ and school performance^48^, lower leisure screen time^49^, and higher chance of smoking cessation^50^. Genomic control inflation in the index GWAS (*γ_GC_* = 1.25; **Supplementary Figure 10**) was explained by polygenicity rather than unadjusted confounding (LDSC intercept = 1.009, se = 0.009). To determine if the index GWAS associations were mostly driven by any single trait, we employed four additional GenomicSEM factor models, each time removing one spousal trait from the analysis. We obtained very high genetic correlations across these analyses (**Supplementary Figure 11**). This suggests that the shared genetic component across spousal traits is robust to the choice of input traits.

We were unable to estimate the heritability of the index (i.e., effective sample size and heritability cannot be simultaneously identified under the factor GWAS framework), but enrichment of heritability based on genomic annotations can be quantified. Heritability of the index showed a 20.2-fold enrichment in conserved DNA elements (p = 9.3e-13; **Supplementary Table 7**) and a 4.5-fold enrichment in functional genomic regions in the anterior caudate (p = 8.0e-4; **Supplementary Table 8** and **Supplementary Figure 12**).

We estimated genetic correlations for the index GWAS with 64 complex traits (**Supplementary Table 9**) and identified many significant correlations (FDR < 0.05; **Figure 3C**; **Supplementary Table 10**). EA showed the highest genetic correlation with the partner choice index (cor = 0.877 and p = 4.3e-460), followed by age at birth of first child (cor = 0.790 and p = 3.2e-236) and household income (cor = 0.761 and p = 2.3e-200). Both the cognitive and non-cognitive components of EA are substantially correlated with the index (cor = 0.524 and 0.701, p = 1.2e-102 and 1.1e-143). Height and BMI are significantly correlated with the index GWAS with opposite signs (cor = 0.197 and -0.399, p = 1.2e-19 and 3.7e-74), while skin and hair pigmentation (within European ancestry) and baldness showed null results. Cigarette smoking and cannabis dependence showed substantial negative correlations with the index GWAS, but drinks per week was not correlated with the index. With respect to personality, neuroticism showed a substantial negative correlation with the index, while openness and agreeableness are positively correlated with the index. We did not find significant correlations for conscientiousness and extraversion. Many health outcomes, including blood cholesterol levels, cardiovascular disease risk, and diabetes were negatively correlated with the index GWAS. Among neurodevelopment and mental health traits, ADHD showed a strong negative correlation with the index GWAS, followed by anxiety, depression, and neuroticism. Anorexia nervosa and bipolar disorder were positively correlated with the index GWAS.

A high genetic correlation between the partner choice index and EA suggests that EA may have a substantial effect on people’s index values in the marriage market. This could be because potential spouses regard those with high EA as more desirable to partner up with, or because those with high EA are able to self-select into environments with a large availability of potential partners who themselves score high on the index. However, an alternative explanation is that the GWAS of EA has substantial uncorrected biases due to assortative mating. Therefore, we obtained summary statistics from within-sibling GWAS analysis^51^ for eight complex traits (**Supplementary Table 11**) and assessed if these sibling GWAS, which are less affected by indirect genetic effects including assortative mating, show similar genetic correlations with the index compared to their population-based GWAS counterparts. We found highly consistent genetic correlations of sibling GWAS with the partner choice index compared to population-based results (**Figure 3D** and **Supplementary Table 12**). In particular, the genetic correlation of the partner choice index with EA remained virtually unchanged when using sibling EA GWAS (cor = 0.871 and se = 0.124). These results suggest that the substantial genetic correlation of EA and the index is not driven by biases in EA GWAS.

### Non-EA component of the partner choice index

Due to the high genetic correlation between the index and EA, we employed GWAS-by-subtraction^52,53^ to search for evidence of other traits contributing to the index that is not explained by EA alone. This approach assumes that genetic associations with partner choice index can be explained by EA genetics and other mechanisms orthogonal to EA (i.e., the non-EA component; **Figure 4A**). It estimates genetic associations with the non-EA component by regressing out the EA genetic associations from index GWAS results.

**Figure 4.**
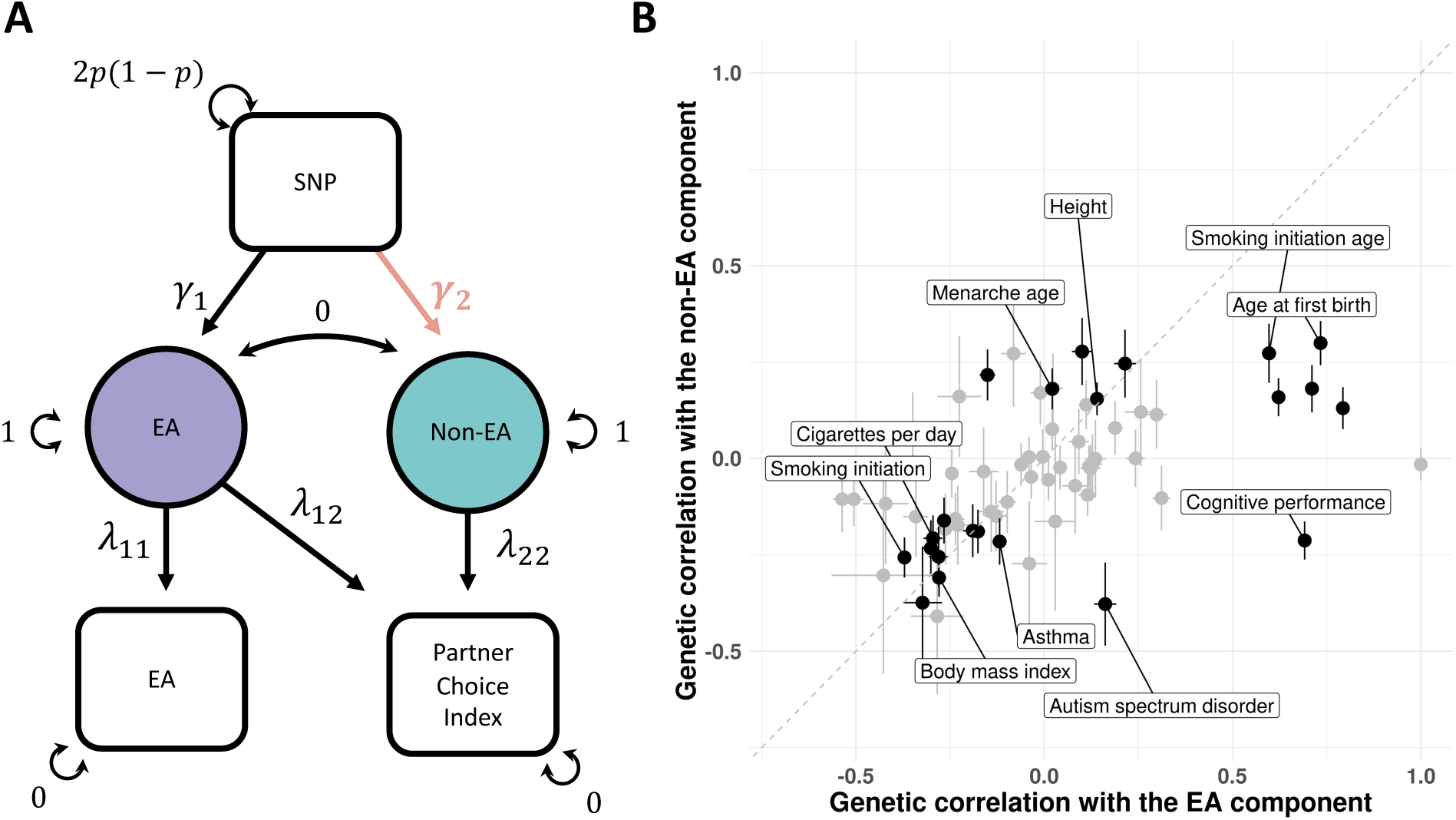
Results of GWAS-by-subtraction. **(A)** Schematic diagram for GWAS-by-subtraction. The main parameter of interest, i.e., genetic effect on the non-EA component *γ*_#_, is highlighted in red. **(B)** Genetic correlations of the EA and non-EA components underlying index GWAS with 64 complex traits. Black data points indicate significant genetic correlations with the non-EA component with FDR < 0.05. Top ten traits showing the most significant correlations with the non-EA component are labeled.

The non-EA component of the partner choice index has a genetic correlation of 0.465 (p = 5.3e-29) with the index itself. We also estimated its genetic correlations with 64 complex traits (**Figure 4B**, **Supplementary Figure 13**, and **Supplementary Table 13**). Height and BMI were both significantly correlated with the non-EA component (cor = 0.155 and -0.309; p = 2.8e-4 and 4.7e-10), showing moderate reductions (∼22%) compared to their correlations with the overall index. Several reproductive and behavioral traits including age at first birth, age at menarche, and cigarette smoking remained significantly correlated but showed substantial reductions. In comparison, asthma showed virtually unchanged genetic correlation with the non-EA component. Autism is a notable trait showing significant correlations with flipped signs with EA (cor = 0.162, p = 4.7e-8) and non-EA components (cor = -0.378, p = 4.6e-4) of the partner choice index, which explains its overall null correlation with index GWAS (cor = -0.022, p = 0.684).

### Comparing GWAS results between sexes and between UK and Norway

Next, we compared genetic associations with the partner choice index between biological sexes and across different geosocial contexts. We performed sex-stratified GWAS meta-analysis on each spousal trait (**Supplementary Table 14** and **Figure 5A**). 4.1% of female BMI variance is explained by male genetics (se = 0.005), which is double the size of how much male BMI is explained by female genetics (h2 = 0.020, se = 0.004). If we assume equal heritability of the male and female indices, this suggests that female BMI has a higher association with the index than male BMI, which is consistent with previous evidence that women’s BMI is more correlated with men’s preference on facial attractiveness than vice versa^54^. In comparison, male EA is more explained by female genetics (h2 = 0.086, se = 0.006) than vice versa (h2 = 0.068, se = 0.006). We employed GenomicSEM single factor models to obtain sex-specific index GWAS (**Supplementary** Figure 14). While some individual spousal traits (e.g. height) showed moderate genetic correlations between sexes, the partner choice index had a near-perfect genetic correlation (**Figure 5B**; cor = 0.974, se = 0.068). They also showed largely consistent genetic correlations with other traits (**Supplementary Table 15** and **Supplementary** Figure 15).

**Figure 5.**
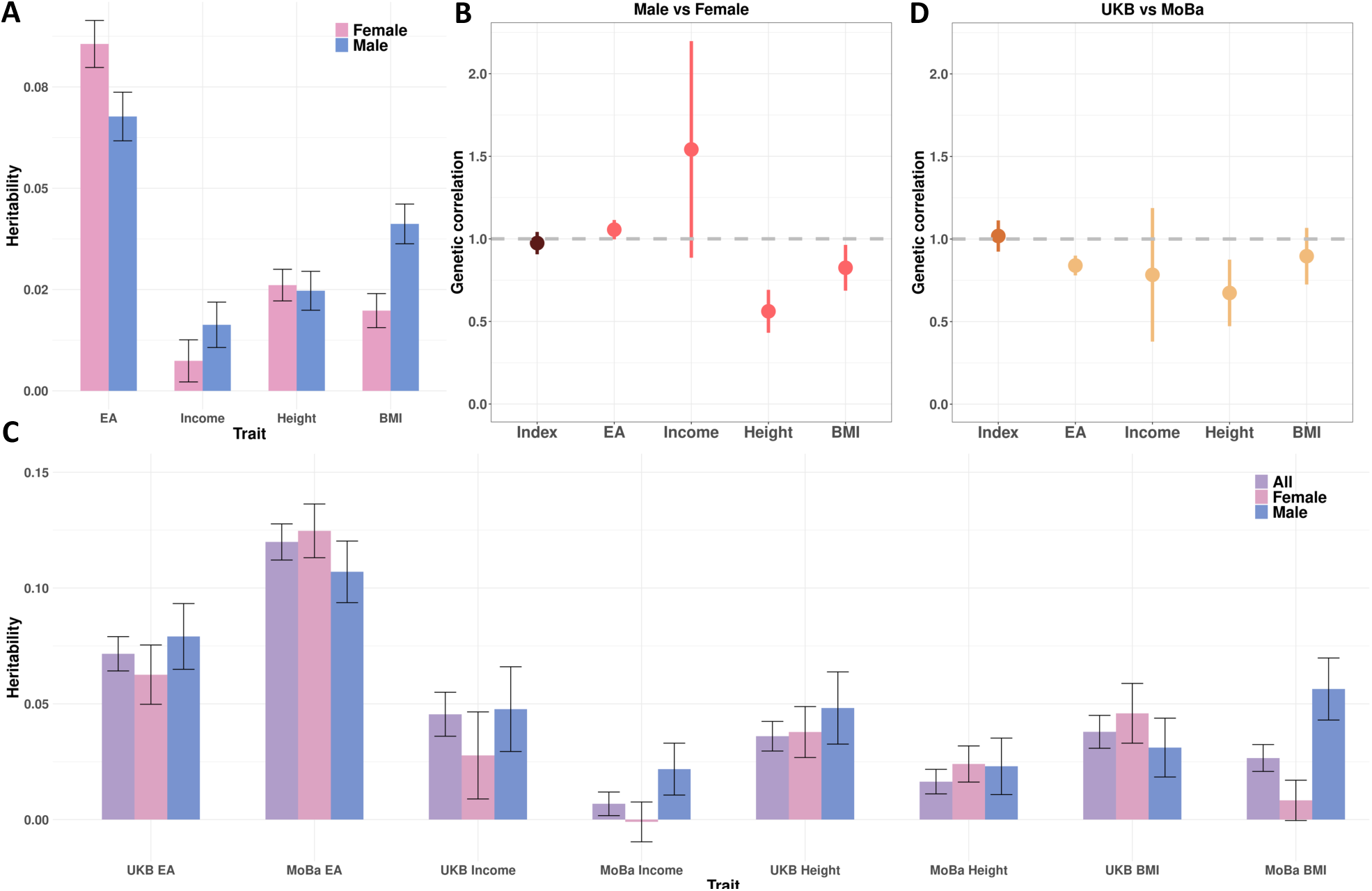
Comparing partner choice indices between sexes and countries. **(A)** Sex-stratified heritability estimates of four spousal traits. Intervals are standard errors. Here, male heritability indicates the fraction of female trait variance explained by male genetics. Similarly, female heritability refers to the fraction of male trait variance explained by female genetics. **(B)** Genetic correlations of partner choice index and four spousal traits between males and females. Intervals are standard errors. **(C)** Heritability estimates of four spousal traits in UKB and MoBa. Intervals are standard errors. **(D)** Genetic correlations of partner choice index and four spousal traits between UKB and MoBa. Intervals are standard errors.

Further, we compared GWAS results between UKB and MoBa, the two cohorts with the largest sample sizes in our study (**Supplementary Table 16**). Spousal EA showed substantially higher heritability in Norway, while heritability of spousal income was higher in the UK (**Figure 5C**). Heritability estimates for spousal height and BMI were both moderately higher in the UK. Sex differences in spousal BMI heritability were much bigger in Norway than in the UK. In MoBa, male heritability of spousal BMI was substantial (h2 = 0.056, se = 0.013) while female heritability was not statistically significant (h2 = 0.008, se = 0.009). We performed GenomicSEM factor GWAS in UKB and MoBa separately (**Supplementary** Figure 16) and obtained a genetic correlation indistinguishable from one (cor = 1.018, se = 0.095), which is higher than the genetic correlations of individual spousal traits (ranging from 0.673 to 0.896; **Figure 5D**). The UKB and MoBa indices also had highly consistent genetic correlations with other complex traits (**Supplementary Table 17** and **Supplementary** Figure 17). We also note that UKB and MoBa participants have different age distributions. Therefore, the heritability differences we observed for individual traits may be explained by generational differences instead of geographic differences. However, the near-perfect genetic correlation of the index throughout these analyses still suggests a highly robust genetic component between sexes, countries, and possibly birth cohorts.

### Removing assortative mating-induced biases in genetic correlation estimates

Next, we investigated whether the pervasive biases^26^ in genetic correlation estimates driven by assortative mating could be remedied by controlling for index GWAS. We focused our analysis on eight complex traits for which summary statistics from both population-based GWAS as well as sibling GWAS were available (**Supplementary Table 11**). For each pair of traits, we estimated genetic correlations using both population-based GWAS and sibling GWAS. Additionally, for both analyses, we produced conditional genetic correlation estimates after controlling for the partner choice index (**Figure 6A**; **Methods**). Full results on these genetic correlations are summarized in **Supplementary** Figure 18 and **Supplementary Table 18**.

**Figure 6.**
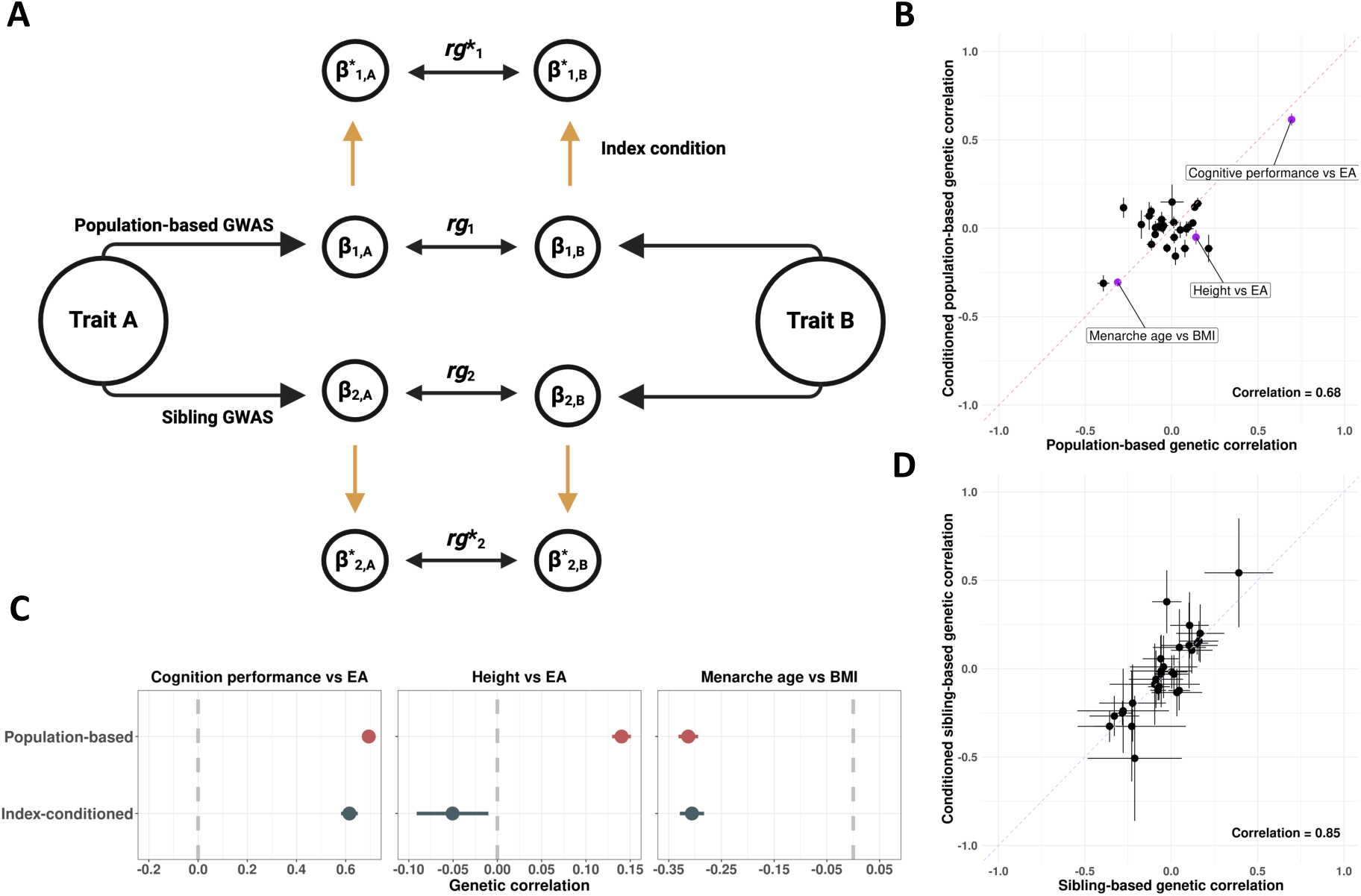
Conditioning on partner choice index in genetic correlation estimation. **(A)** Comparative schematic of four sets of genetic correlations. We estimated regular and index-conditioned genetic correlations across eight complex traits using both population-based and sibling GWAS data. Details on calculation of conditional genetic correlation are shown in **Methods** and **Supplementary** Figure 21. **(B)** Unconditioned and index-conditioned genetic correlations estimated from population-based GWAS of eight complex traits. Three trait pairs mentioned in the main text are highlighted. Error bars show standard errors. **(C)** Population-based genetic correlations between cognitive performance and EA, height and EA, and menarche age and BMI. Error bars are standard errors. **(D)** Unconditioned and index-conditioned genetic correlations estimated from sibling GWAS of eight complex traits. Error bars show ±1 standard errors.

Conditioning on the partner choice index substantially reduced genetic correlations between many trait pairs (**Figure 6B**). For example, the significant and likely spurious correlation between height and EA (cor = 0.14, p = 8e-40) was reduced to null after conditioning on the index (cor = -0.05, p = 0.21; **Figure 6C**). Importantly, some genetic correlations were not altered. Examples include the correlations between menarche age and BMI and between EA and cognitive performance (**Figure 6C**), suggesting that these population-based estimates were not severely biased due to uncorrected assortative mating in GWAS. Across trait pairs, index-conditioned and unconditioned estimates had a moderate correlation of 0.68. In comparison, sibling GWAS are known to be robust to indirect genetic mechanisms including assortative mating^51,55,56^. Indeed, conditioning on the index did not substantially alter genetic correlation estimates obtained from sibling GWAS (**Figure 6D**). Despite imprecision in these sibling-based correlation estimates due to lower GWAS sample size, index-conditioned and unconditioned estimates were highly concordant (correlation = 0.85), showcasing the validity of our approach. An additional attractive feature of our correction approach is that index-conditioned genetic correlations are more precisely estimated than the sibling-based genetic correlations. Based on sample sizes in this analysis, index-conditioned population-based analysis led to an average 62% reduction in the size of standard errors across trait pairs.

### PGS-level footprint of assortative mating and out-of-sample prediction in UKB

Finally, given the well-powered index GWAS, we can now produce PGS for the partner choice index. We explored two applications of this PGS using data from UKB participants of European ancestry who have not been included in the GWAS (N = 320,142). First, we investigated the long-range LD patterns consistent with index-driven assortative mating using the correlation of PGS derived from odd and even chromosomes^22^ (**Methods**). We found a highly significant odd-even chromosomal correlation (cor = 0.025, p = 1.1e-44, **Supplementary Table 19**). Although we also found similar correlations for PGS based on GWAS of height, BMI, EA, and income, these correlations were substantially attenuated after conditioning on the partner choice index (**Figure 7**), which again suggests that the partner choice index GWAS can be used to correct GWAS statistics for assortative mating (**Methods**). By contrast, height remained statistically significant after conditioning on the index.

**Figure 7.**
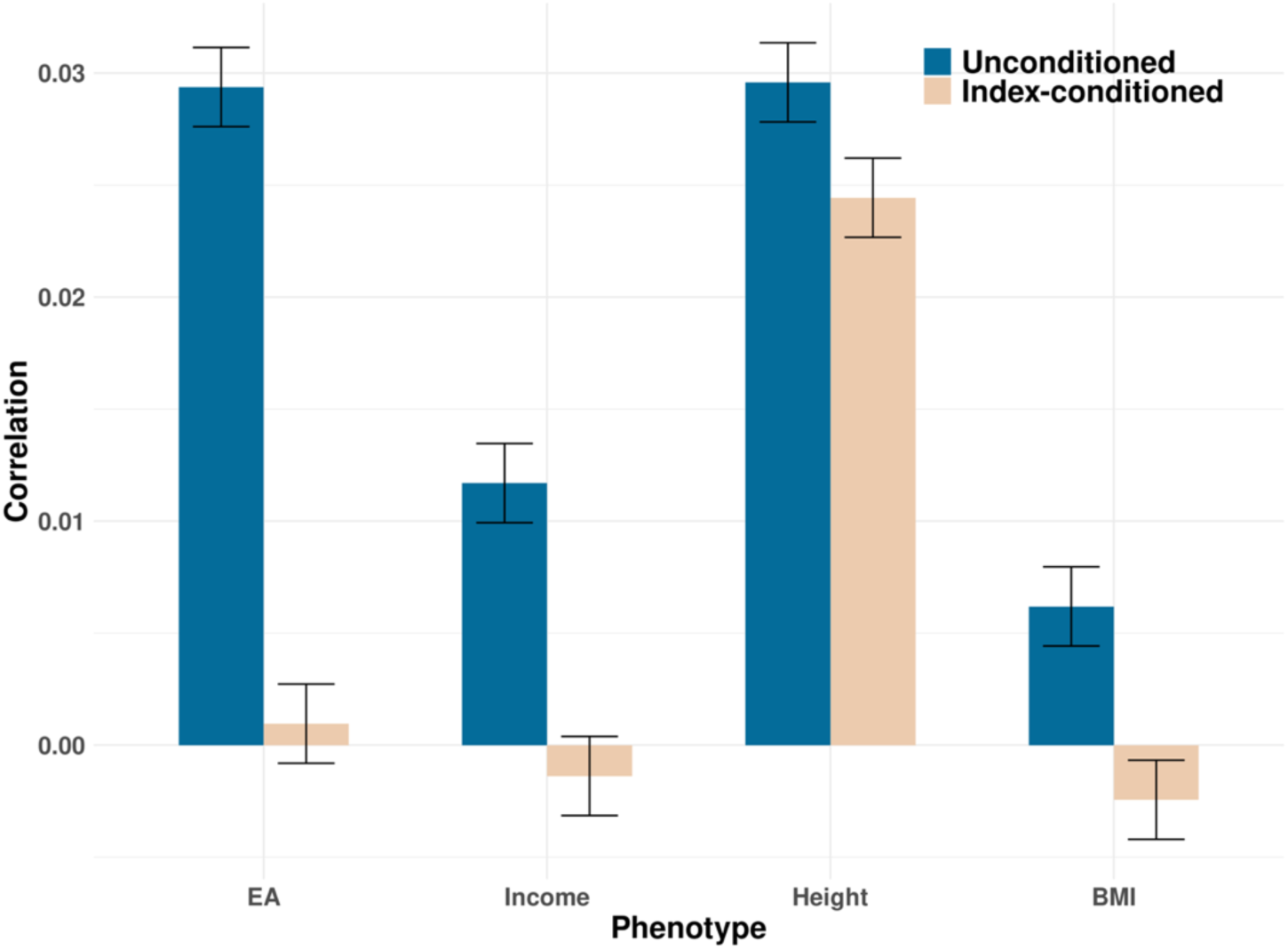
Correlation between PGS based on odd and even chromosomes. Unconditioned correlations were obtained using PGS derived from population-based GWAS. Index-conditioned results were based on residual GWAS after controlling for partner choice index. Error bars are ±1 standard errors.

Next, we tested for associations of the partner choice index PGS based on the full SNP data of all chromosomes with (own) EA, income, BMI, and height in UKB. We found highly significant associations between the index PGS and all traits (**Supplementary Table 20**). The incremental R^2^ after controlling for sex, age, and genetic principal components were low, ranging from 0.39% to 2.02%. We also calculated PGS using the sex-specific index GWAS and found differences in their predictive performance in male and female samples. The Male index PGS was more associated with height, while the female index PGS was more strongly associated with lower BMI (**Supplementary** Figure 19). These results provide further evidence that BMI is more associated with women’s index than in men. In comparison, height may be correlated more strongly with men’s index. Finally, we performed PGS association analyses using sibling samples in the UKB (**Methods**).

Associations of the index PGS with various phenotypes were substantially attenuated in within-sibling analysis compared to estimates obtained from independent individuals (on average, regression coefficients reduced by 62.8% across traits), but association with EA and income remained highly significant (**Supplementary** Figure 20 and **Supplementary Table 21**).

## Discussion

In human societies, individuals match to form couples, start families, and have children. Human genetics is both a cause and a product of this matching process, leading to well-documented phenotypic and genetic patterns of non-random mating. Here we present the first ever common factor GWAS on partner characteristics, contributing to a deeper understanding of partner preferences and resulting assortative mating patterns. Using four genotyped global cohorts, we find genetic associations and substantial SNP-based heritability for four spousal characteristics: EA, income, height, and BMI. High genetic correlations amongst the four spousal characteristics are consistent with our approach’s assumption that partners match along a single index, an important characteristic of several theoretical economic models of the marriage market^34^. However, we also made important observations that these genetic correlations are not exactly 1 (or -1), indicating that both matching on a latent index and trait-specific sorting might play a role. Based on this evidence, we designed and employed a factor GWAS strategy to identify a common factor across four spousal characteristics. We demonstrated that a GWAS of this common factor robustly identifies genetic associations with the partner choice index even when the single index assumption is violated. Leveraging these methodological advances, this study is the first to probe deeper into the mechanisms through which assortative mating occurs (i.e., empirically identifying the latent index driving assortative mating), and how this leaves imprints in the human genome (i.e., obtaining genetic associations for the index).

Our analyses revealed substantial genetic correlations between the partner choice index and various complex traits, which sheds important light on the numerous characteristics that contribute to assortative mating. In particular, we found a substantial genetic correlation of the index with EA, but GWAS-by-subtraction analysis indicates that genetic correlations between the index and many complex traits do not operate through EA alone. Importantly, these genetic correlations also suggest that SNP associations with many traits may be partially reflecting uncorrected, assortative-mating-driven biases in existing GWAS. While such biases may be weak for individual SNPs, they could have substantial cumulative impact on numerous genetic epidemiologic applications involving genome-wide SNP data. Based on this insight, we designed an index-conditioned analytical strategy to control for assortative mating in (1) PGS correlations between odd-even chromosomes and (2) genetic correlations between complex traits. We demonstrated that our index-conditioned genetic correlation analysis produces similar and statistically more efficient estimates compared to the gold-standard sibling-based estimates. Most notably, the positive and significant genetic correlation between EA and height, previously reported in the literature^26,51,57^, is reduced to zero after conditioning on the partner choice index.

The genetic associations between spousal characteristics and the partner choice index could arise from various mechanisms that may be complex and dependent on social structure. Potential mechanisms include an individual’s preference for partner characteristics, but also effects that may operate through social and cultural phenomena such as the institutions, networks and social arenas in which individuals interact with others^58–61^. For example, the finding that SNPs associated with EA also correlate with partner’s EA could arise because college-educated individuals are more likely to meet their partner in environments where college graduates are overrepresented, or express their personal preference in college-educated individuals for finding a college-educated partner. Further, the strength of SNP associations may depend on cultural and social norms regarding partner choice. Hence, assortative mating patterns we have uncovered are a result of both choice and opportunity structure. Alternatively, indirect genetic effects such as uncontrolled population stratification may drive GWAS signal: SNPs that correlate with geographic differences may be associated with the partner choice index, especially when these geographic differences are positively correlated with socioeconomic status^62,63^. We note that patterns of homogamy and heterogamy in couples are influenced not only by the mating process, but also by later demographic events, such as union dissolutions and re-partnering. It is important to bear in mind the complex social mechanisms behind these associations when interpreting our findings.

Our study has several limitations. First, emerging studies suggest evidence of a multi-index sorting framework^64^. Our framework has extended beyond a single-index model by allowing trait-specific matching (i.e., individuals prefer partners who are similar to themselves, such that SNPs associated with trait X correlate with trait X in one’s spouse; **Supplementary** Figure 1**, Supplementary Note**). In our analysis, spousal height exhibits patterns consistent with both index-driven and trait-specific matching, showing a lower genetic correlation with other spousal traits while its odd-even chromosomal PGS correlation remains significant, albeit attenuated, after controlling for the index GWAS. The use of single-index GWAS summary statistics to correct GWAS results for assortative mating will remove bias due to index-driven assortative mating, but bias due to trait-specific assortative mating may remain. A full investigation of multi-index sorting and their genetic associations requires GWAS on a broader collection of spousal characteristics and likely larger study samples. Second, sample selection is a limitation. Our sample is limited to individuals clustering with European ancestry reference populations, as there is a dearth of genetic data on couples of other ancestries^65^. The focus on individuals who are coupled excludes those who are single. We also focused exclusively on different-sex couples, which was necessary to accurately impute spouses in the UKB. Importantly, the issue of non-random sample selection also contributes to the heterogeneity across study cohorts. We combined data from cohorts with relatively wide ranges in age which may mask major changes in preferences for partner characteristics in response to societal changes. The MoBa sample is further limited by including only couples who have had a common child. The families where the father also participates in MoBa are known to be more selected than families where the father is not participating. Similarly, UKB is not representative of the general UK population^66^ and couples who both participate in UKB are likely to be more selected than the rest of UKB. These issues, coupled with other sources of heterogeneity such as different phenotypic definitions across cohorts (e.g., we used occupational income in the UKB but observed annual income reported by tax authorities in MoBa), may reduce the replicability and statistical power of our results. However, we were encouraged by (a) the genome-wide significant loci identified in GWAS meta-analysis, which suggests consistent SNP-level associations across cohorts, and (b) the near-perfect genetic correlations of the partner choice index between sexes, and between UKB and MoBa, which suggests a surprisingly robust single index driving assortative mating in highly variable geosocial contexts. Still, due to these limitations, an important future direction is to expand the study cohort by including couples from diverse ancestries and cultures, and perform within-family GWAS to further tease apart direct genetic effects on the index from other mechanisms leading to indirect associations. Third, we had limited consideration in this study on how assortative mating patterns differ between generations and whether it has reached equilibrium in the population^19^. Finally, although the index PGS may have applications in the future (beyond predicting the four traits we included in this study), we caution that the predictive performance of the current PGS, although statistically significant, is not yet practically meaningful.

In conclusion, our results show that GWAS associations of various spousal characteristics, combined with economic methods, increase our knowledge about the genetic architecture of assortative mating. It also highlights the importance of studying the social structures that give rise to assortative mating, as well as the genetics of the underlying latent traits that matter for the matching of partners. Studying the genetic imprints of assortative mating can help correct for bias in various GWAS-related statistics, and further our understanding of the causes and consequences of assortative mating, and its implications for social structures and human behavior.

## Methods

### Cohort data

We imputed spousal pairs in UKB following a recent study^18^. Couples were identified based on the following criteria: report living with their spouse (data field 6141), report the same length of time living in the house (699), report the same number of occupants (709) and number of vehicles (728) in the household, report the same accommodation type and rental status (670, 680), have identical home coordinates (rounded to the nearest km) (20074, 20075), and are registered with the same UKB recruitment center (54). We require both individuals to have genotype data and answer above questions. In results, we obtained 39,169 spousal pairs of European ancestry in UKB. Genetic ancestry was obtained from UKB data field 22006. Of the four phenotypic characteristics, height is obtained from data field 12144 and BMI is calculated using height and weight (data field 21002). We calculated log hourly wage following Kweon et al.^67^ and years of education using data field 6138, following Lee et al.^68^.

HRS is a biannual and nationally representative longitudinal survey of older adults in the United States. We restrict the analytical sample to different-sex couples of European ancestry in HRS. Respondents who can be linked with two or more different partners via the spousal ID variable were omitted from the analysis. Spouses who have the same self-reported sex were also excluded. These restrictions led to a sample of 14,396 couples.

MoBa is a population-based pregnancy cohort study conducted by the Norwegian Institute of Public Health.^69^ Participants were recruited from all over Norway from 1999-2008. Women consented to participation in 41% of the pregnancies. Blood samples were obtained from both parents during pregnancy and from mothers and children (umbilical cord) at birth^70^. The cohort includes approximately 114,500 children, 95,200 mothers and 75,200 fathers. Genotyping, quality control, imputation, and post-imputation of the MoBa genotype data were made available through the MoBaPyschGen project^71^. The current study is based on version 12 of the quality-assured survey data files released for research in January 2019. The establishment of MoBa and initial data collection was based on a license from the Norwegian Data Protection Agency and approval from The Regional Committees for Medical and Health Research Ethics. The MoBa cohort is currently regulated by the Norwegian Health Registry Act. The University of Oslo data protection official (PVO) has approved the data protection impact assessment (DPIA) for this project. An agreement with MoBa allows access and use of MoBa data. In this study, MoBa was linked to the National Educational Database and the Income and Taxation register, which contains information on EA and annual income, respectively, for the whole Norwegian population. EA information was taken from registers for the age range 35-50 for the year that is closest to the year of birth of the MoBa child. Income information was taken from the two years prior to the birth of the MoBa child. Height and BMI data was reported in the MoBa questionnaire data (at 15 weeks of pregnancy) supplemented with information from the Medical Birth Register of Norway. Missing paternal data was filled in using maternal reports from the MoBa questionnaire. Couples were identified from MoBa recruitment and genotype data (N = 77,012).

The Lifelines Cohort Study is a multi-disciplinary prospective population-based cohort study examining the health and health-related behaviors of 167,729 persons living in the North of the Netherlands in a three-generation design^72^. Between 2006 and 2013, randomly selected general practitioners invited all their listed patients aged 25-49 years to participate in the study. Subsequently, these participants invited their family members to join, such that individuals from all age ranges were included. Currently, around 80,000 of the 167,729 respondents have been genotyped. Partner status was self-reported by the respondents, and we only included observations for which both the respondent and the partner were participants in Lifelines. We dropped individuals younger than 30 years, such that our Lifelines GWAS sample consisted of 17,702 respondents. 8,630 were male and 9,072 were female. As phenotypes, we used partner’s years of education, partner’s height and partner’s BMI. Years of education was constructed based on the highest degree obtained as reported by the respondent. Height and BMI were measured during the first assessment visit. We did not use income, as the only measure of income available in Lifelines was household income.

Additional details of phenotypic definition in each cohort are provided in **Supplementary Table 22**.

### GWAS implementation

Four traits were included in the analysis: height, BMI, income, and EA. For each trait, we performed GWAS on both own and spousal outcomes using individuals whose spousal data were available in each cohort. GWAS were conducted using PLINK2.0^73^ while adjusting for sex, age, genotyping array (in UKB), and the top 20 principal components. We removed SNPs with a missing call rate > 0.01, a minor allele frequency (MAF) < 0.01, and a Hardy–Weinberg equilibrium test P-value < 1e-6 from the analyses. Additionally, to explore sex-specific genetic associations, we also conducted sex-specific GWAS in each cohort while dropping sex as a covariate. We performed fixed-effect meta-analysis using METAL^74^ to combine associations for each trait across four cohorts. Due to the lack of individual-level income data in Lifelines, meta-analysis for income and spousal income included data solely from the UKB, HRS, and MoBa.

We derived partner choice index GWAS using GenomicSEM^43^. To obtain the genetic covariance (S) matrix and corresponding sampling covariance matrix (V), we ran multivariable LDSC on HapMap3 SNPs, using 1000 Genomes Project European samples as the LD reference. For quality control, we excluded SNPs with a MAF < 0.05 in factor GWAS. Four sensitivity analyses were carried out for factor GWAS. Each time, one of the four spousal trait GWAS was excluded, while the remaining three were used to fit the single factor model and obtain genetic associations for the index.

We looked up the within-family genetic associations for the genome-wide significant loci identified in spousal trait GWAS meta-analysis. We identified 19,136 full sibling pairs among UKB participants of European ancestry using KING^75^. We further employed SNIPar^56^ to impute the sum of parental genotypes of these sibling pairs. We then extracted 5,701 unrelated individuals from the intersection of sibling pairs and spousal pairs in UKB to form the sample for within-family association analysis. We used SNIPar to conduct association analysis on spousal phenotypes with parental genotype added as covariates.

We also performed sex-specific and cohort-specific index GWAS. We performed meta-analyses of sex-specific GWAS, and then used the summary statistics as input to obtain sex-specific index GWAS associations. Cohort-specific analysis focused exclusively on UKB and MoBa due to the larger sample size of these two biobank cohorts.

We used LDSC^76^ to perform heritability and genetic correlation analysis, using the 1000 Genomes Project European samples as the LD reference. We deployed heritability enrichment analysis using stratified LDSC^77^ with the baselineLD V2.2^78^ genomic annotations and GenoSkyline-Plus^79^ tissue and cell-type annotations. We employed GSUB^52^, a close-form approach to perform GWAS-by-subtraction, to obtain the non-EA component GWAS underlying partner choice index. EA summary statistics in the GWAS-by-subtraction analysis was obtained from the largest EA GWAS to date^80^.

### PGS analysis

We used GWAS summary statistics for the partner choice index to calculate PGS for UKB samples of European ancestry. We included independent samples (N = 320,142) that were unrelated to the 39,169 spousal pairs used in the GWAS analysis.

We applied PUMAS-EN^81^, an elastic net ensemble learning algorithm using 4-fold Monte Carlo cross-validation (MCCV) that adaptively selects and aggregates many input PGS models to generate SNP weights from GWAS summary statistics. Within each fold of MCCV, we partitioned full GWAS summary statistics into PGS training (70%) and ensemble training (30%) summary statistics. We included PGS models lassosum^82^, LDpred2^83^, PRS-CS^84^, MegaPRS^85^, and SDPR^86^ with tuning parameter settings previously described^81^. We used 1000 Genomes Project European samples as the LD reference and HapMap 3 SNPs for all PGS models and ensemble training.

To estimate the correlation between PGS derived from odd and even chromosomes, we calculated two sets of PGS for each individual – one using SNPs located on odd chromosomes and the other using SNPs on even chromosomes. We computed Pearson correlation between these two sets of PGS to quantify evidence for long-range LD and assortative mating. In the index-conditioned analysis, we first used GSUB to obtain the residual non-index-explained genetic component for each trait (**Supplementary** Figure 21). Then, we used the residual GWAS (i.e., non-index component) for each trait to compute PGS and odd-even-chromosomal correlations.

In the PGS-phenotype association analysis, we created PGS using genome-wide SNP data. These scores were standardized to a mean of 0 and variance of 1. We regressed each standardized phenotype on the index PGS, while controlling for sex, age at recruitment, and 20 principal components. We performed both sex-combined and sex-specific analyses. PGS generated from sex-specific index GWAS were used in sex-specific PGS-phenotype association analysis. For within-sibling analysis, we identified UKB full-sibling pairs (from different families) using KING^75^. 11,391 sibling pairs from different families that are also unrelated to our UKB GWAS discovery sample were included in the analysis. We assessed the association between delta-phenotype (i.e., difference of standardized phenotype between siblings) and delta-PGS (i.e., difference of index PGS between siblings) after controlling for the sibling difference of covariates.

### Estimation of conditional genetic correlation

Consider traits *Y*_1_, *Y*_2_, and *Z* whose additive genetic components we denote as *G*_Y_1__, *G*_/Y2_, and *G_Z_*, respectively. We define the conditional genetic correlation between traits *Y*_1_ and *Y*_2_, given trait *Z*, as the correlation of *G*_Y_1__ and *G*_Y_2__ given *G_Z_*. This can be estimated using *cor*(*δ*_Y_1__, *δ*_Y_2__), where *δ*_Y_1__ (*δ*_Y_2__) is the residual of *G*_Y_1__ (*G*_Y_2__) after projecting onto *G*_0_ . To estimate *cor*(*δ*_Y_1__, *δ*_Y_2__), we first apply GSUB^52^ to obtain residual GWAS of *Y*_1_and *Y*_2_ after subtracting GWAS of *Z*, which essentially provides GWAS summary statistics for *δ*_Y_1__ and *δ*_Y_1__. Then, we apply bivariate LDSC to residual GWAS summary statistics to estimate the conditional genetic correlation *cor*(*δ*_Y_1__, *δ*_Y_2__). In our application, we used index GWAS as trait *Z* to adjust for biases in the genetic correlation estimates for other trait pairs.

## Data availability

GWAS summary statistics produced in this study are available at https://qlu-lab.org/data.html

## Acknowledgments

We thank David Hugh-Jones, Matthew Keller, and members of the Social Genomics Working Group at University of Wisconsin for helpful discussions. We are grateful for the feedback we have received at The Advances in Social Genomics Conference, American Society of Human Genetics Annual Meeting, Integrating Genetics and the Social Sciences Conference, and Center for Economic and Social Research at University of Southern California when an early version of this work was presented. This research has been conducted using the UK Biobank Resource under Application 42148. The Norwegian Mother, Father and Child Cohort Study is supported by the Norwegian Ministry of Health and Care Services and the Ministry of Education and Research. We are grateful to all the participating families in Norway who take part in this on-going cohort study. We thank the Norwegian Institute of Public Health (NIPH) for generating high-quality genomic data. This research is part of the HARVEST collaboration, supported by the Research Council of Norway (RCN; #229624). We also thank the NORMENT Centre for providing genotype data, funded by the Research Council of Norway (HSØ; #223273), South-Eastern Norway Health Authorities and Stiftelsen Kristian Gerhard Jebsen. We further thank the Center for Diabetes Research, the University of Bergen for providing genotype data and performing quality control and imputation of the data funded by the ERC AdG project SELECTionPREDISPOSED, Stiftelsen Kristian Gerhard Jebsen, Trond Mohn Foundation, the Research Council of Norway, the Novo Nordisk Foundation, the University of Bergen, and the Western Norway Health Authorities. The Moba analyses were on the Tjeneste for Sensitive Data (TSD) facilities, owned by the University of Oslo, operated and developed by the TSD service group at the University of Oslo, IT-Department (USIT), using resources provided by Sigma2—the National Infrastructure for High Performance Computing and Data Storage in Norway (UNINETT). ECC is supported by HSØ (#2021045) and RCN (#274611). SA was supported by the European Union’s Horizon 2020 research and innovation programme under the Marie Skłodowska-Curie grant agreement (ESSGN 101073237), the National Institute on Aging of the National Institutes of Health (R01AG078522, and R01AG079554), and the Dutch National Science Foundation (016.VIDI.185.044). The Lifelines initiative has been made possible by subsidy from the Dutch Ministry of Health, Welfare and Sport, the Dutch Ministry of Economic Affairs, the University Medical Center Groningen (UMCG), Groningen University and the Provinces in the North of the Netherlands (Drenthe, Friesland, Groningen). The authors wish to acknowledge the services of the Lifelines Cohort Study, the contributing research centres delivering data to Lifelines, and all the study participants. AH is supported by the RCN (#274611, #288083, #336085, #300668), Helse Sør-Øst (#2020022, #2019097, ##2021045) and the European Union’s Horizon 2020 Research and Innovation program (FAMILY, #101057529; and Marie Skłodowska-Curie grant ESSGN #101073237). The Health and Retirement Study (HRS) genetic data were accessed through NIAGADS with accession number NG00119.v1. These data were collected with financial support from the National Institute of Health’s (NIH) Director’s Opportunity for Research awards using American Reinvestment and Recovery Act funds (RC2 AG036495-01, RC4 AG039029-01). With these funds, the HRS has genotyped almost 20,000 respondents who provided DNA samples and signed consent forms in 2006–2012. The HRS data were produced and distributed by the University of Michigan under the directorship of David R. Weir, with funding from the National Institute on Aging (grant number NIA U01AG009470), Ann Arbor, MI. Figure 1 was created in BioRender: Zheng, Q. (2025) https://BioRender.com/v48y355.

## Author contribution

Q.L. conceived and designed the study.

Q.Z. performed statistical analyses.

S.A. performed GWAS in Lifelines.

T.H.L. performed GWAS in MoBa.

Q.L., E.C., and P.A.C. derived the theoretical framework for index-based sorting.

Z.S., J.M., Y.W., and S.D. assisted with post-GWAS analyses.

B.Z. advised on spouse imputation.

M.N. advised on GenomicSEM.

P.T. advised on assortative mating-driven biases in post-GWAS applications.

J.F., A.H., E.C.C., and T.J.G advised on study cohorts.

J.F. advised on social science issues.

Q.L., Q.Z., S.A., and T.H.L wrote the manuscript. All authors revised and approved the manuscript.

