## Supplementary Figures for "Genetic basis of partner choice"

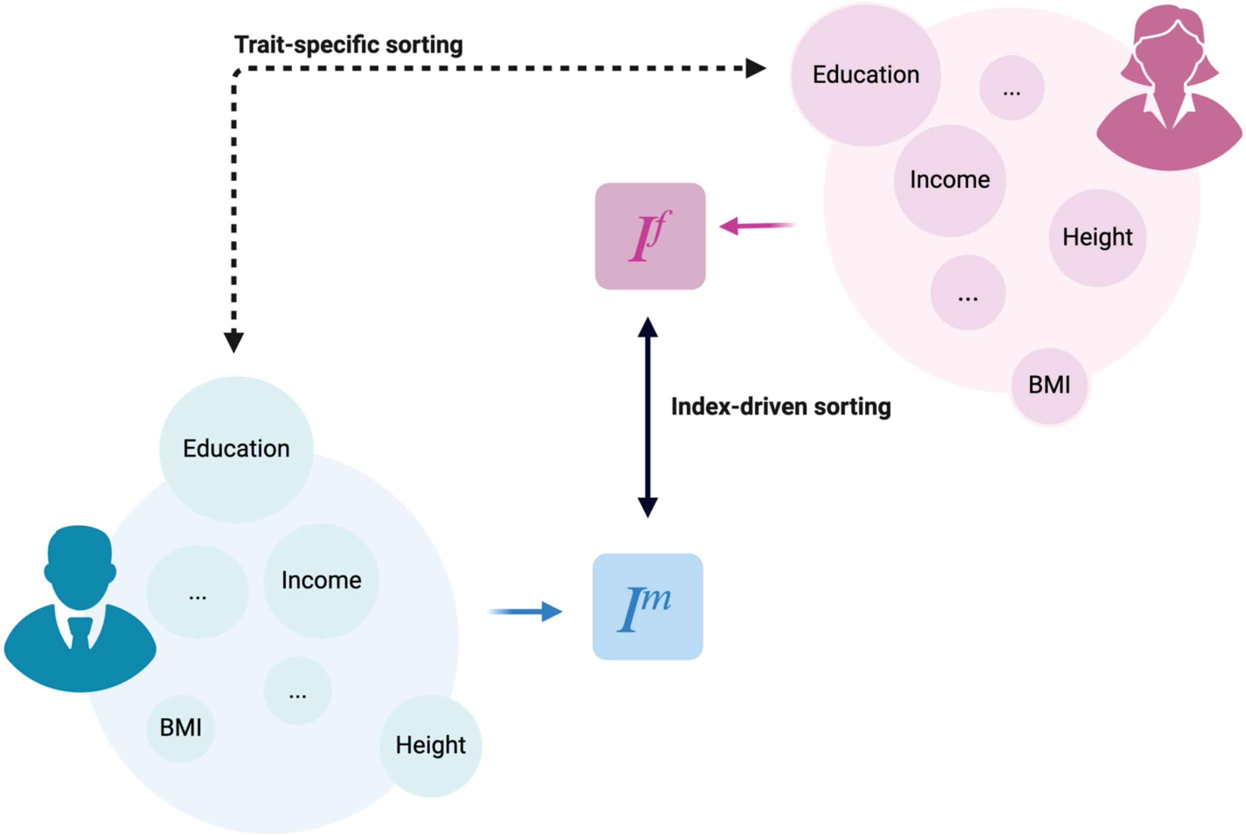


**Supplementary Figure 1. A framework allowing partner choice to be driven by both index-based and trait-specific sorting mechanisms.**

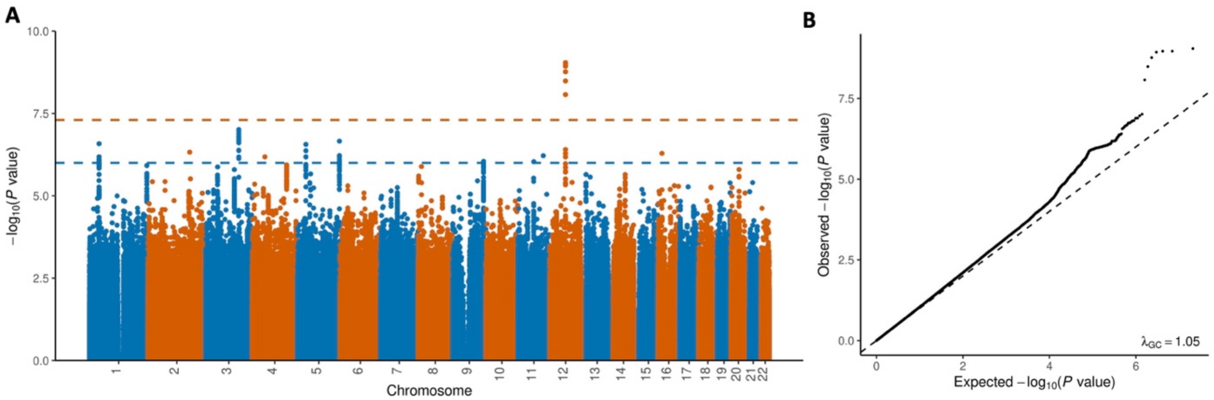


**Supplementary Figure 2. A) Manhattan plot and B) QQ plot for the GWAS meta-analysis on spousal height.**

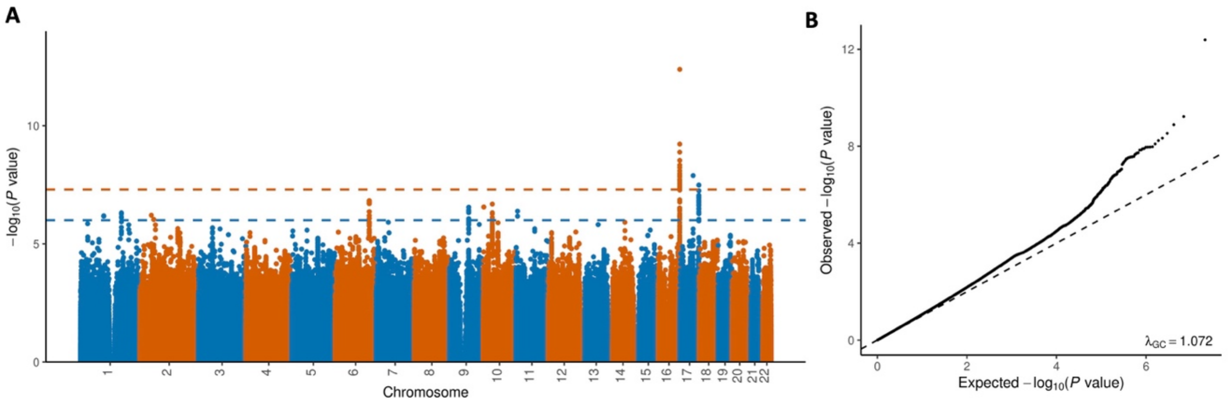


**Supplementary Figure 3. A) Manhattan plot and B) QQ plot for the GWAS meta-analysis on spousal BMI.**

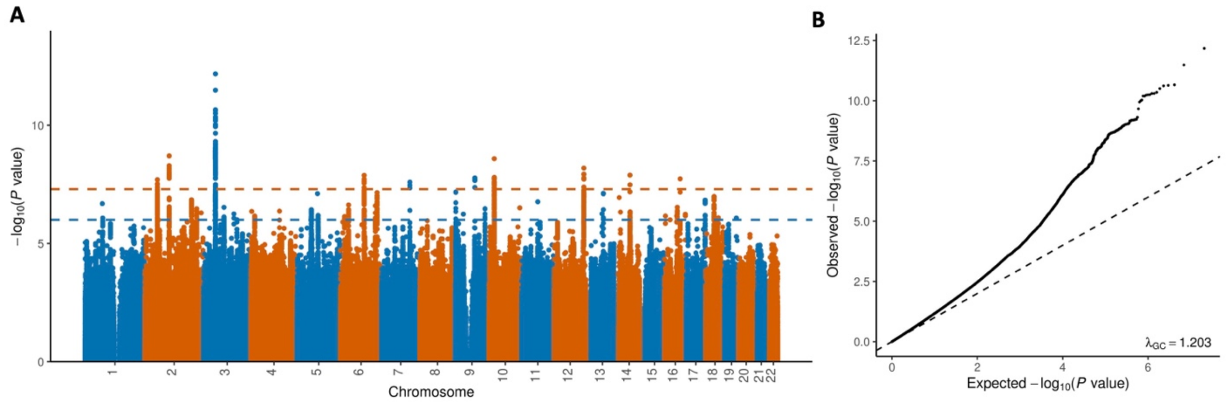


**Supplementary Figure 4. A) Manhattan plot and B) QQ plot for the GWAS meta-analysis on spousal educational attainment.**

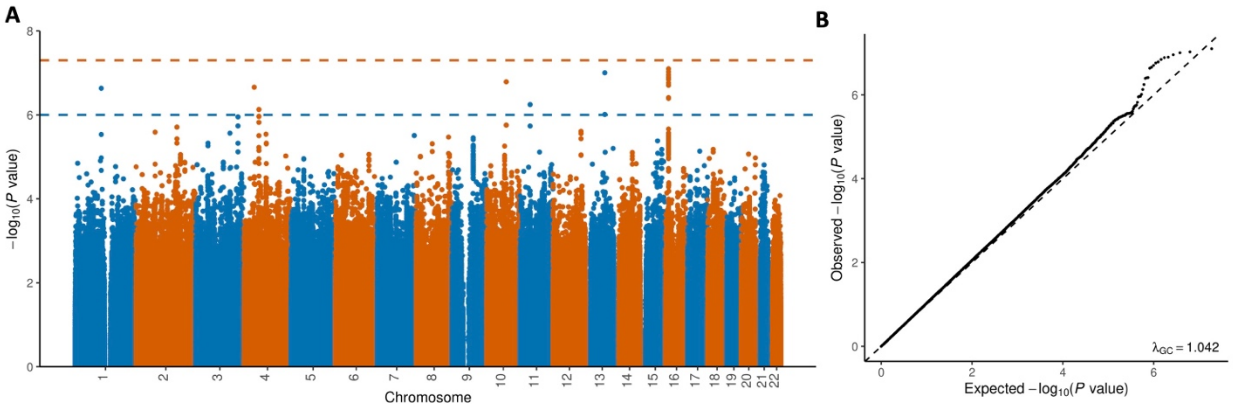


**Supplementary Figure 5. A) Manhattan plot and B) QQ plot for the GWAS meta-analysis on spousal income.**

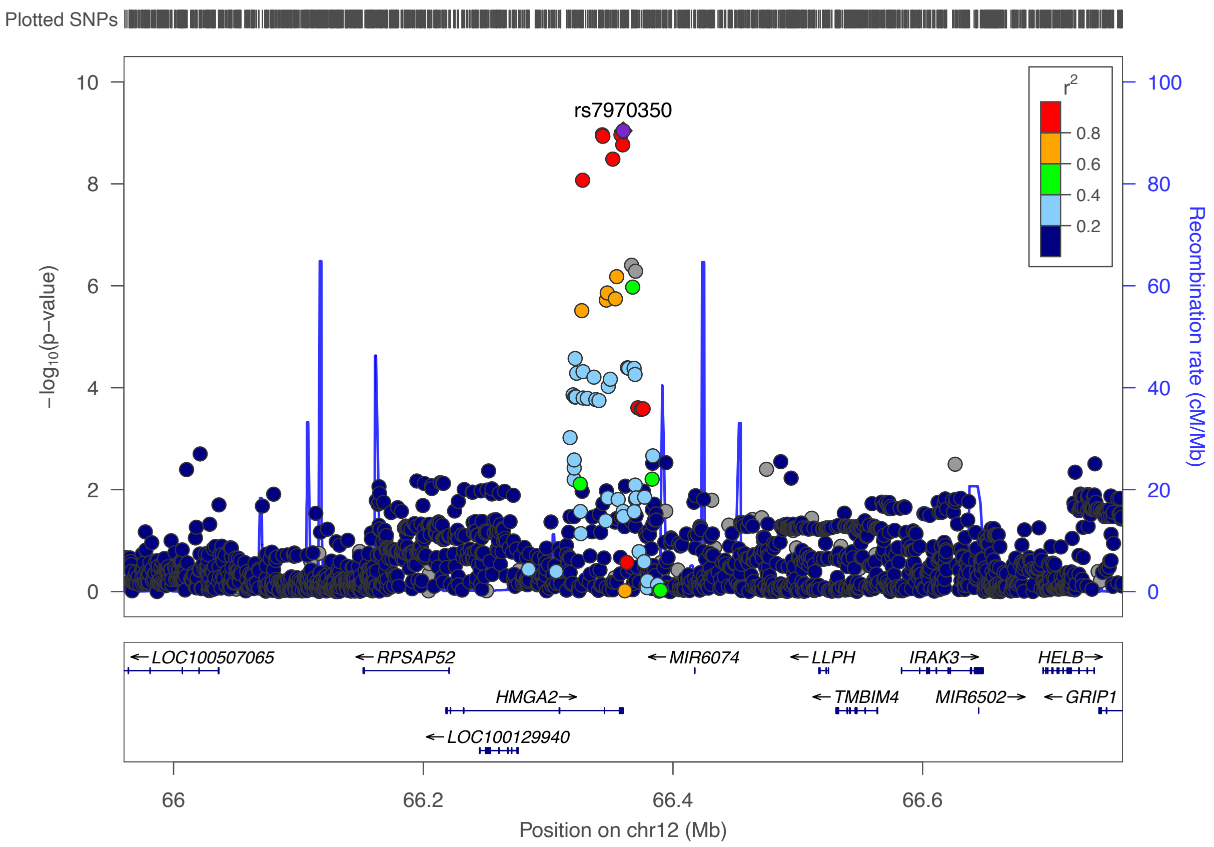


**Supplementary Figure 6. Genome-wide significant association from the GWAS on spousal height.**

**A**

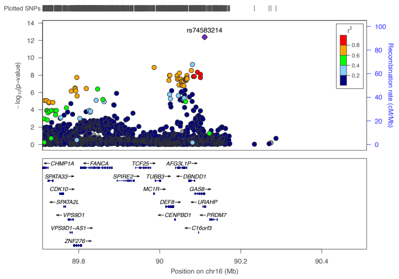


rs74583214

**B**

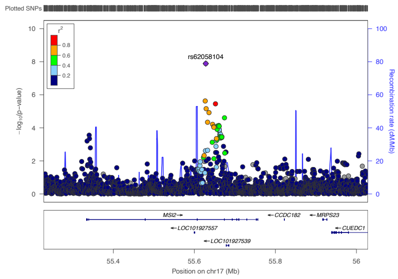


rs62058104

**C**

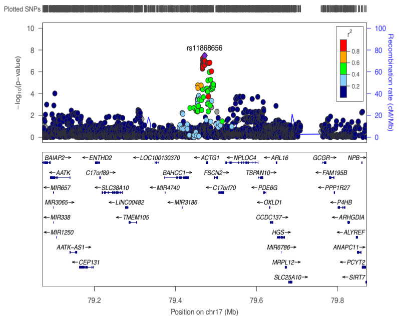


rs11868656

**Supplementary Figure 7. Genome-wide significant associations from the GWAS on spousal BMI.**

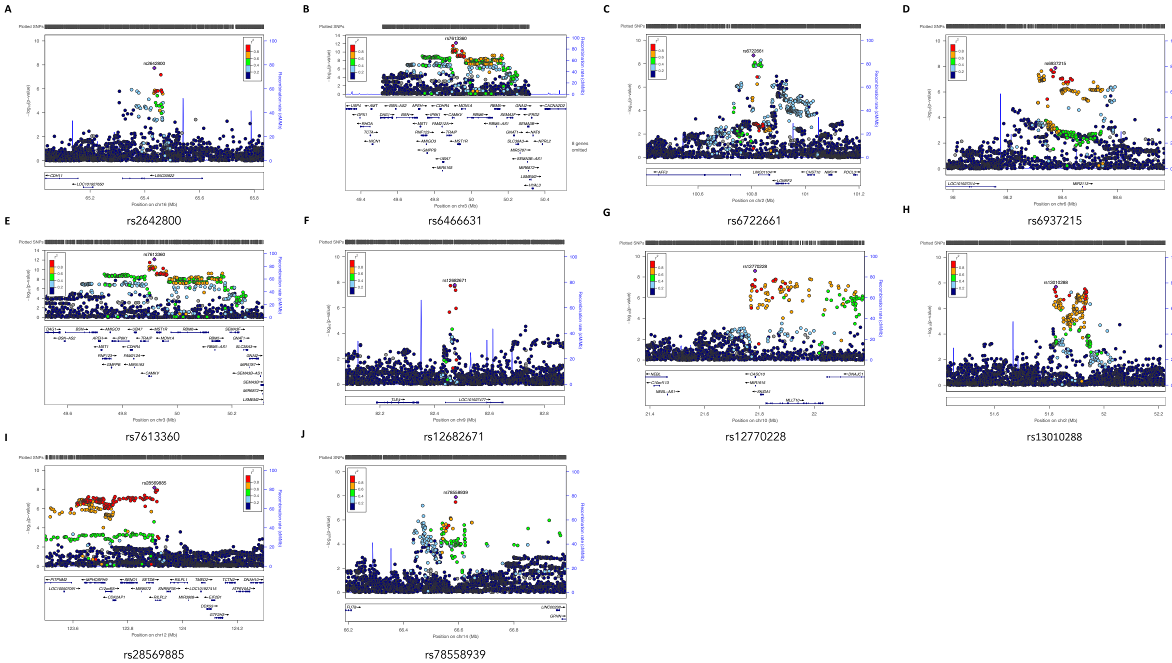


**Supplementary Figure 8. Genome-wide significant association from the GWAS on spousal education attainment.**

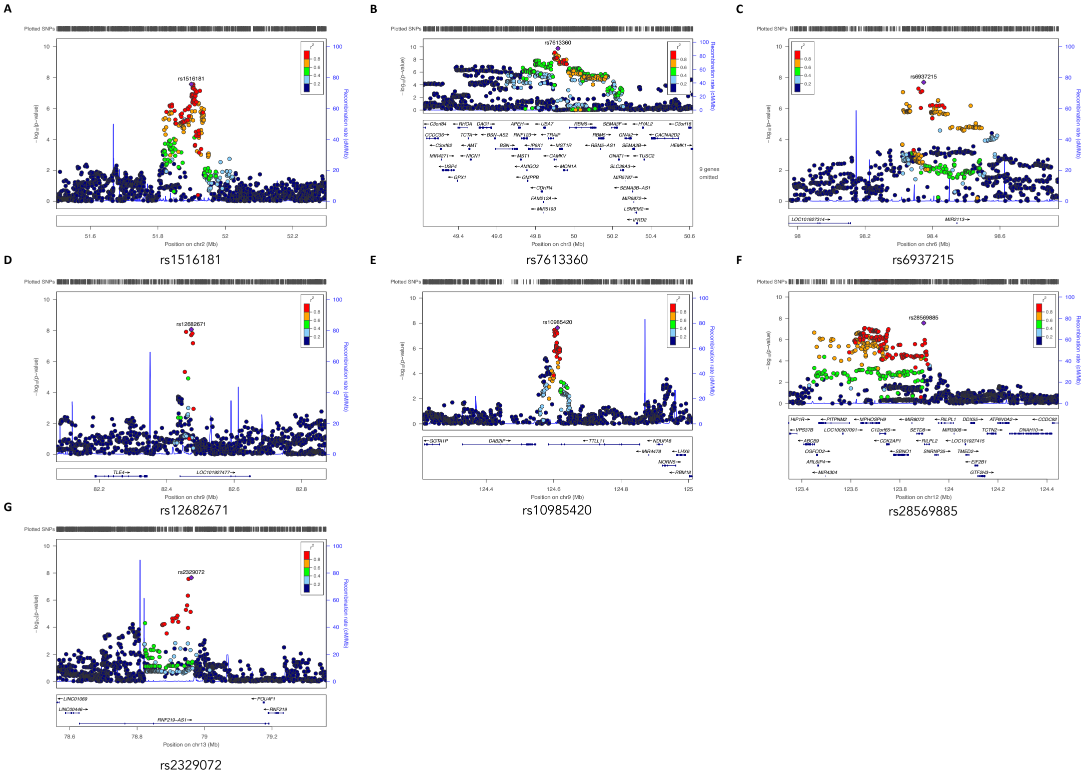


**Supplementary Figure 9. Genome-wide significant associations from the factor GWAS based on four spousal trait GWAS meta-analysis.**

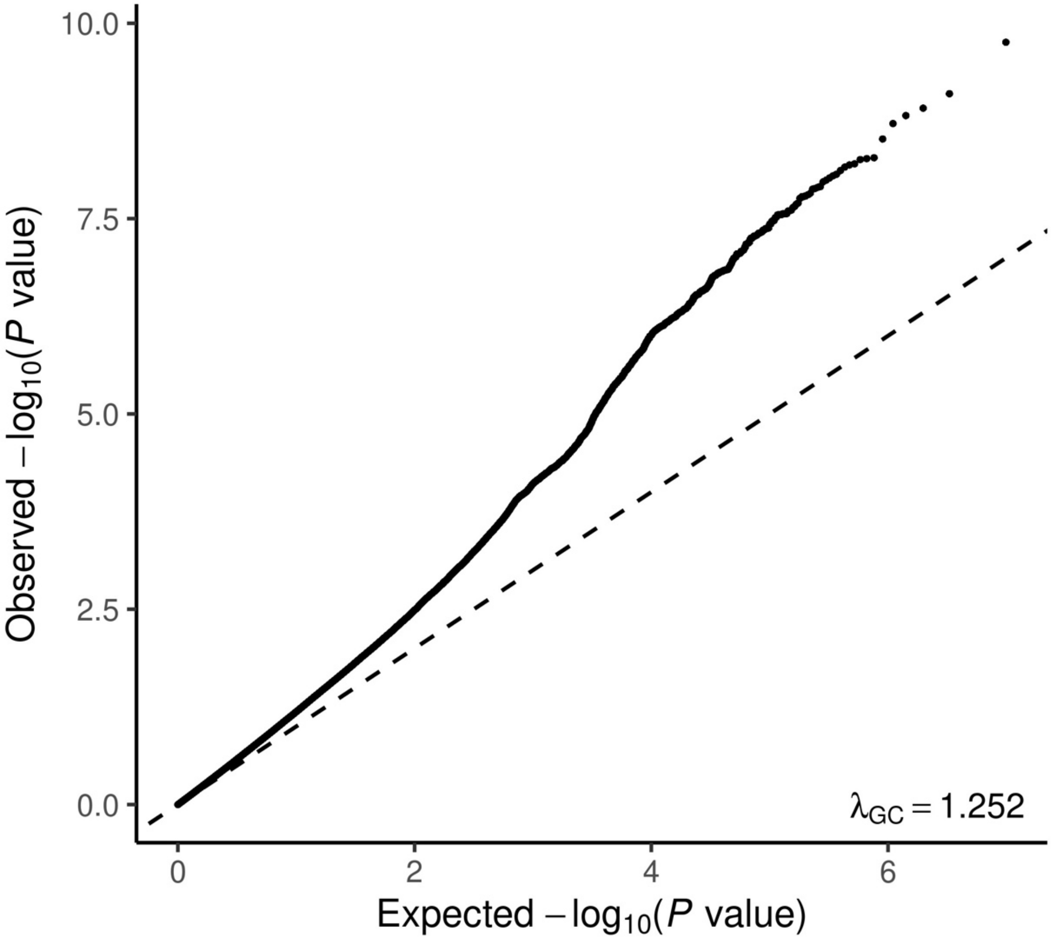


**Supplementary Figure 10. QQ plot for the factor GWAS based on four spousal trait GWAS meta-analysis.**

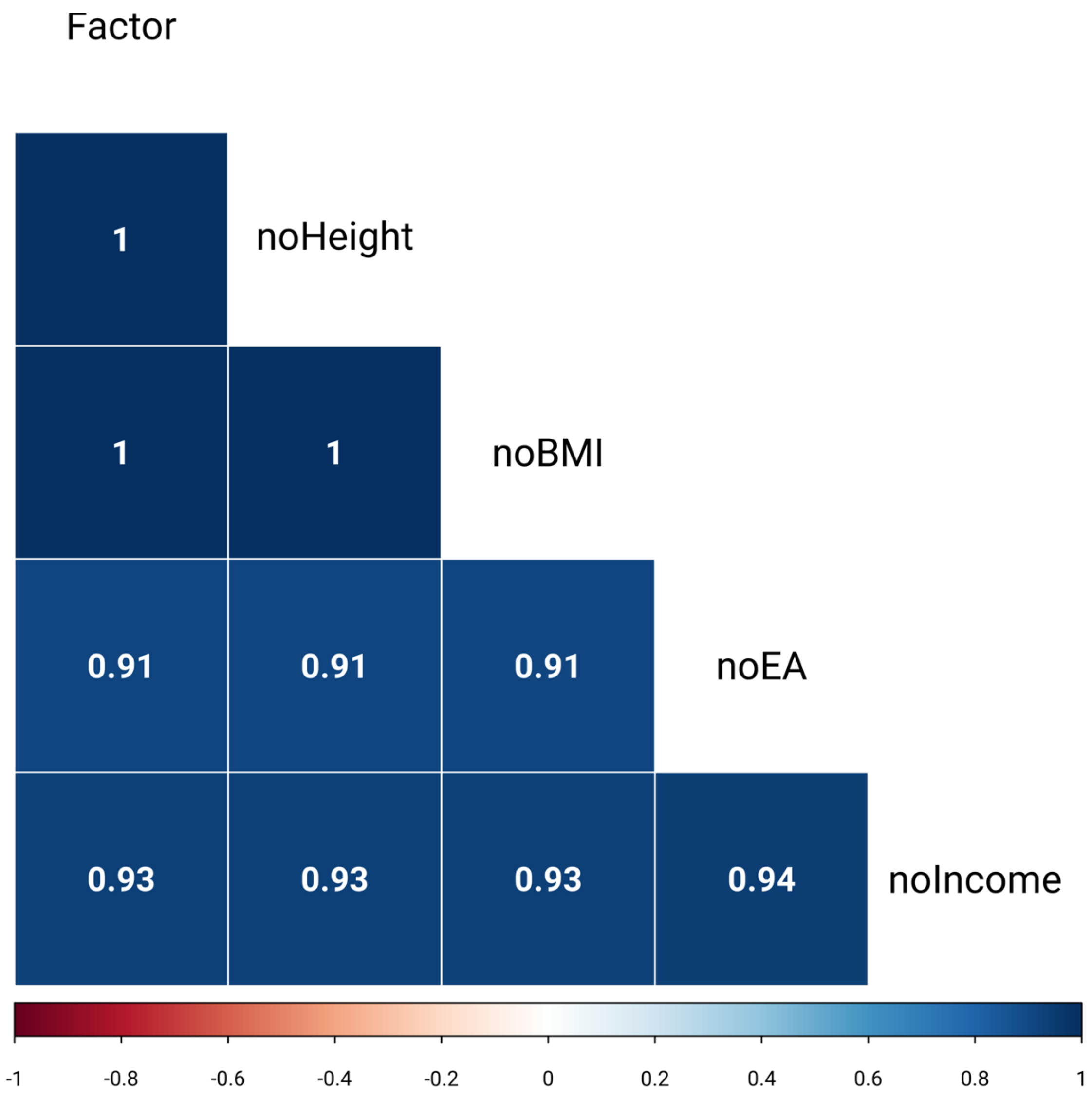


**Supplementary Figure 11. Genetic correlations between the four-trait factor GWAS and three-trait factor GWAS.**

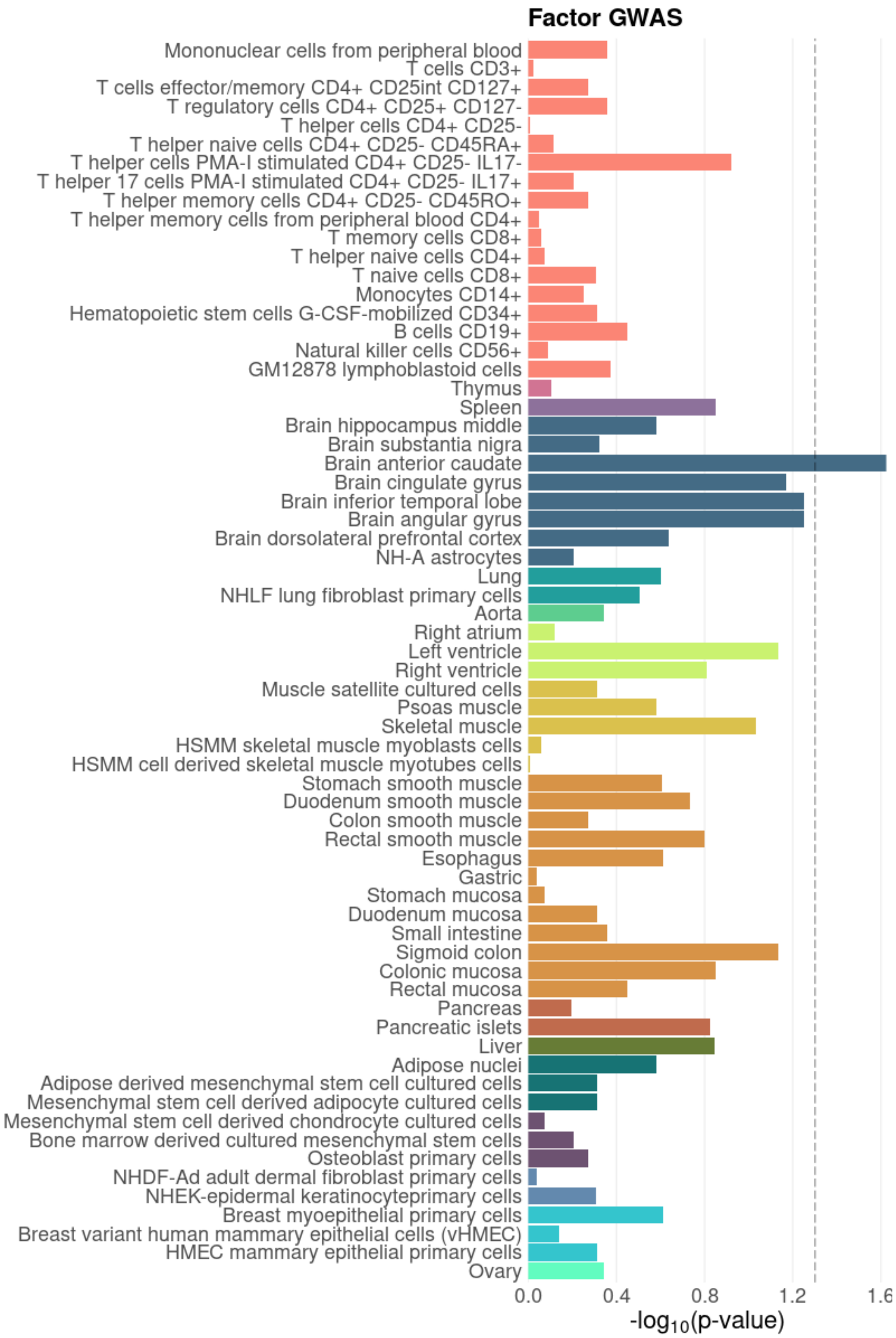


**Supplementary Figure 12. Heritability enrichment of the factor GWAS in functional genomic regions in 66 tissue and cell types predicted by GenoSkyline-Plus.** Dotted line indicates the FDR-corrected significance threshold.

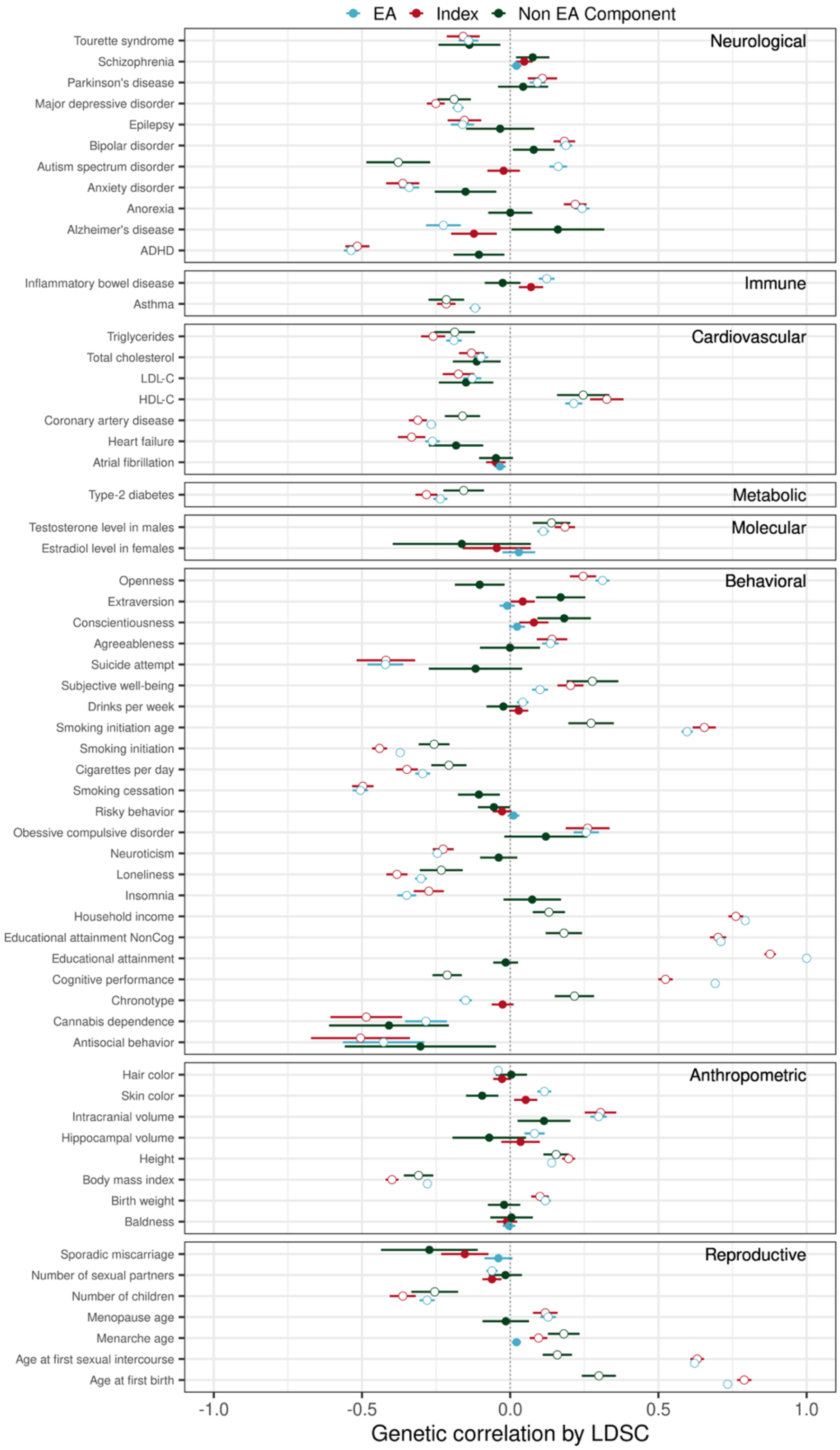


**Supplementary Figure 13. Genetic correlations of the EA and non-EA components underlying partner choice index with 64 complex traits.** Genetic correlations of the index with other traits are also shown as a reference. Hollow circles indicate significant correlations at FDR < 0.05. Intervals indicate standard errors.

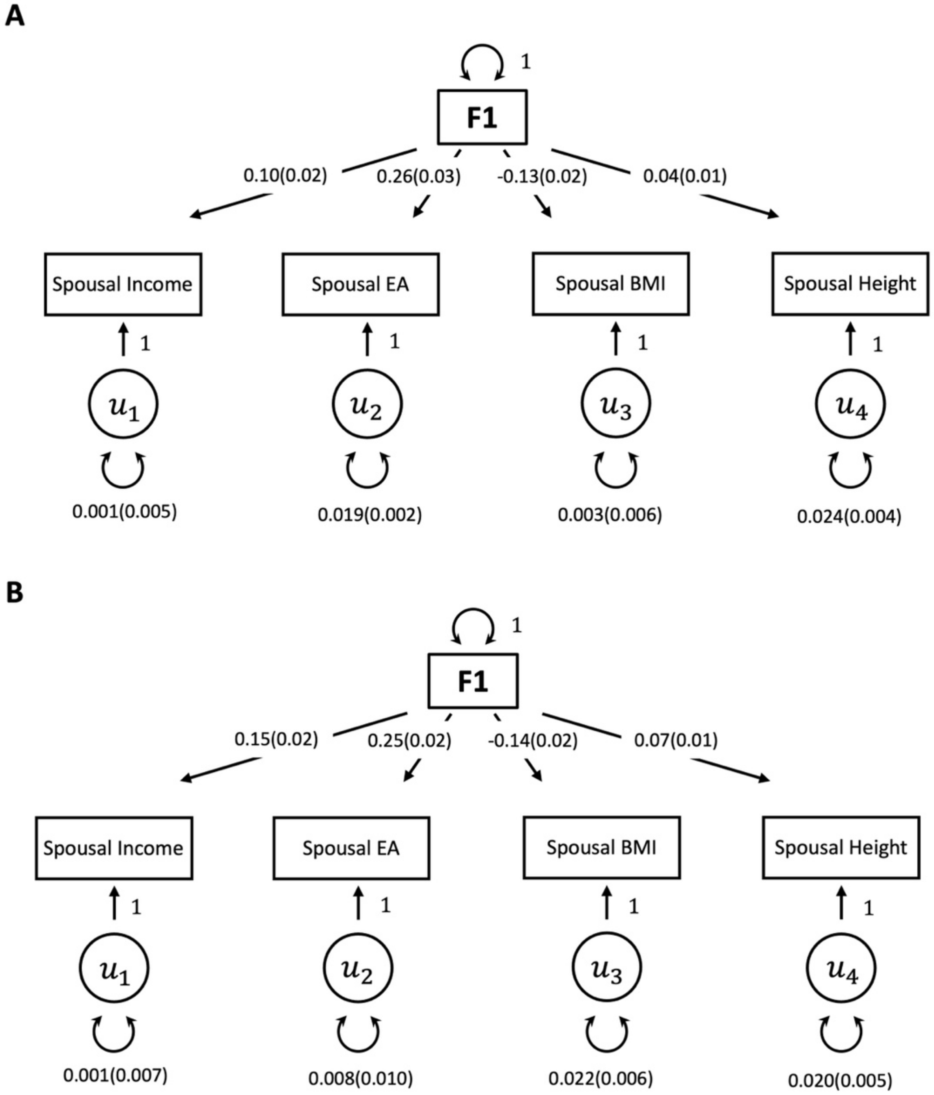


**Supplementary Figure 14. GenomicSEM factor model fit using A) female-specific GWAS and B) male-specific GWAS.**

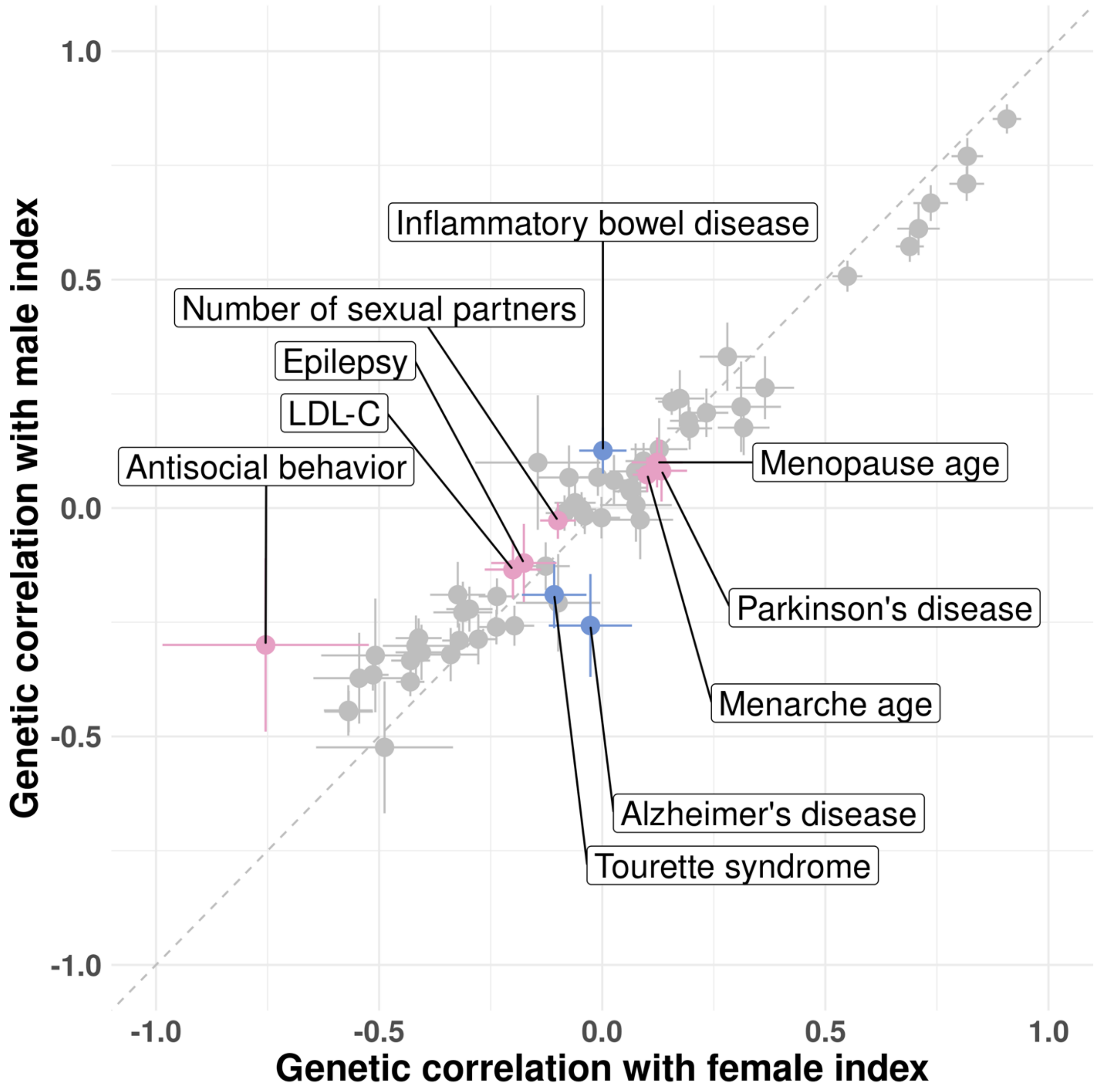


**Supplementary Figure 15. Genetic correlations of male and female indices with 64 complex traits.** Highlighted data points indicate traits showing significant correlations (FDR < 0.05) with only the male or the female index. Intervals indicate standard errors.

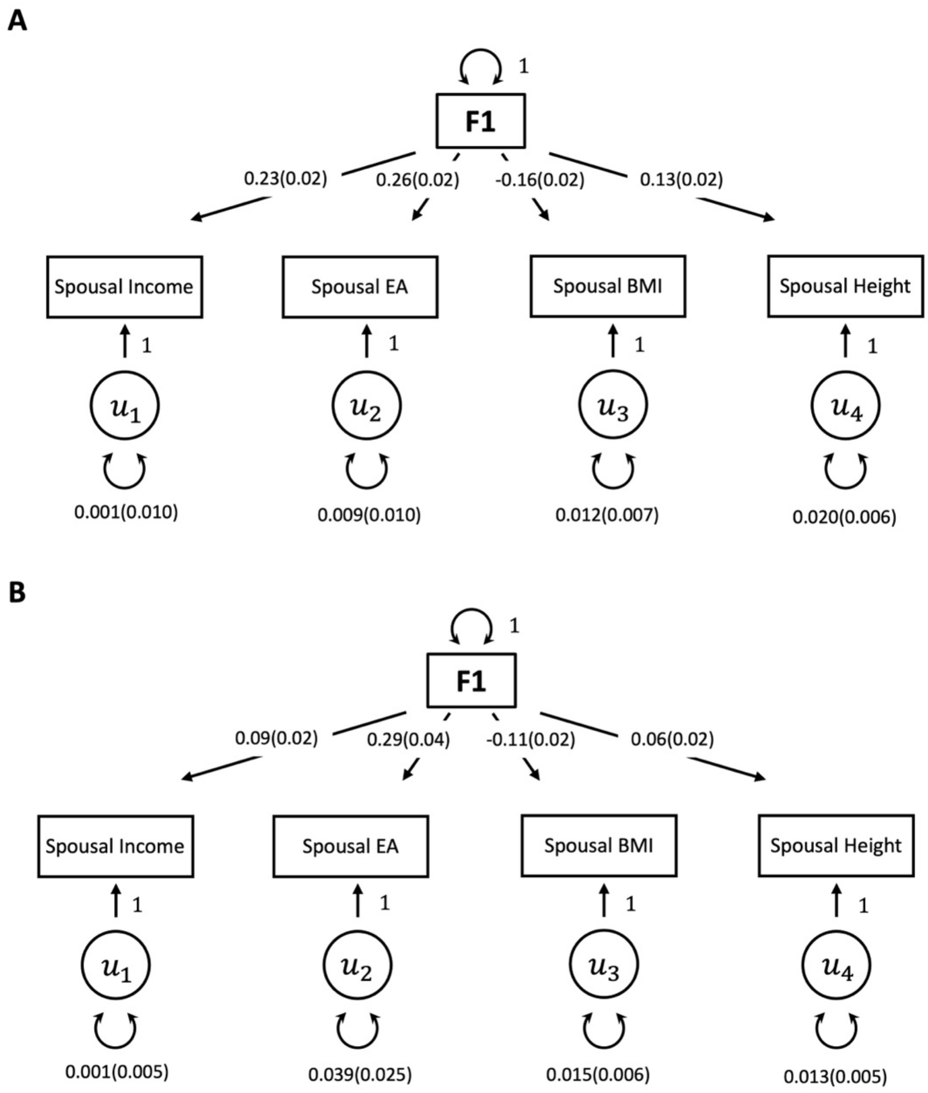


**Supplementary Figure 16. GenomicSEM factor model fit in A) UKB and B) MoBa.**

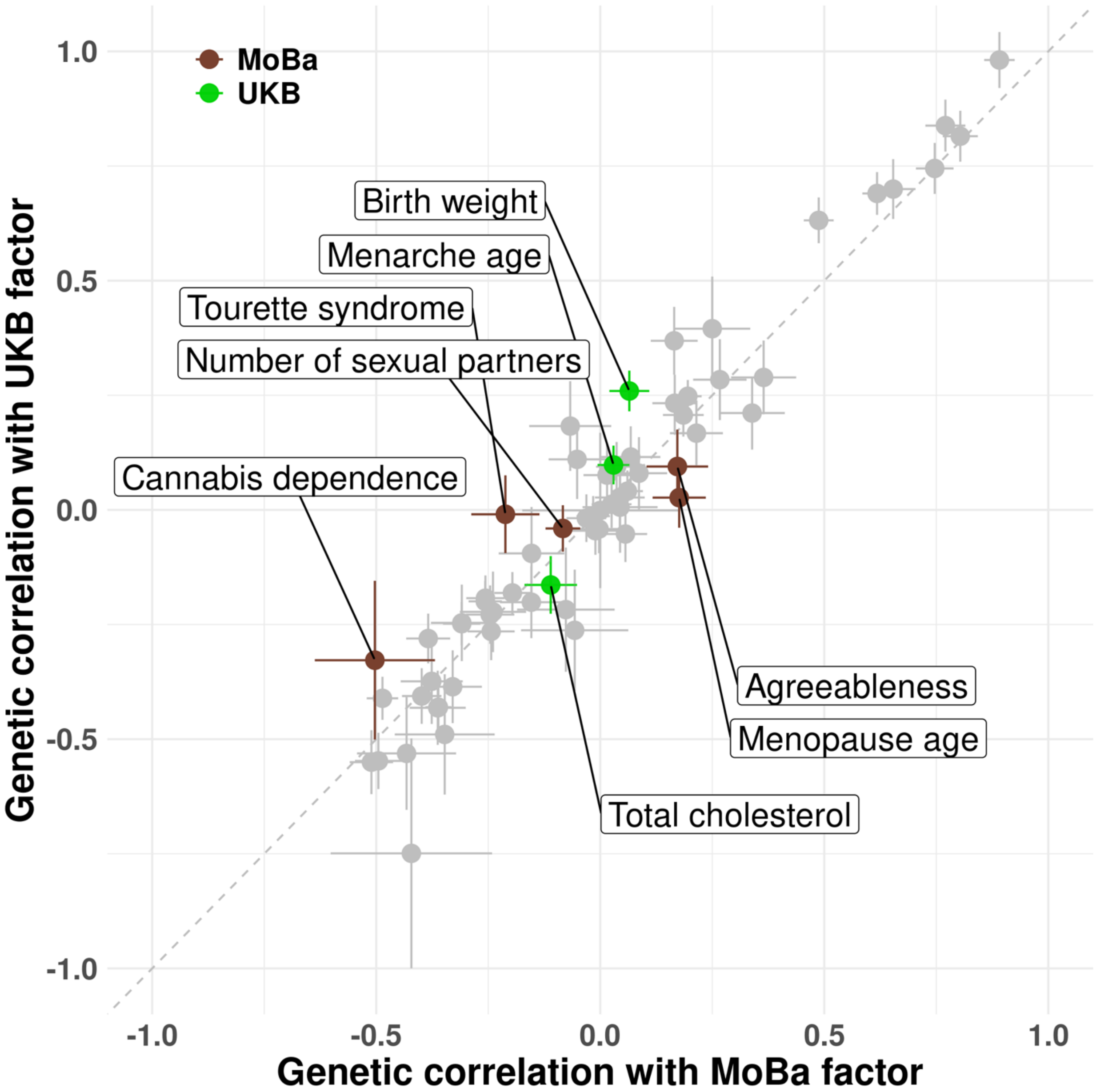


**Supplementary Figure 17. Genetic correlations of UKB and MoBa indices with 64 complex traits.** Highlighted data points indicate traits showing significant correlations (FDR < 0.05) with only the UKB or the MoBa index. Intervals indicate standard errors.

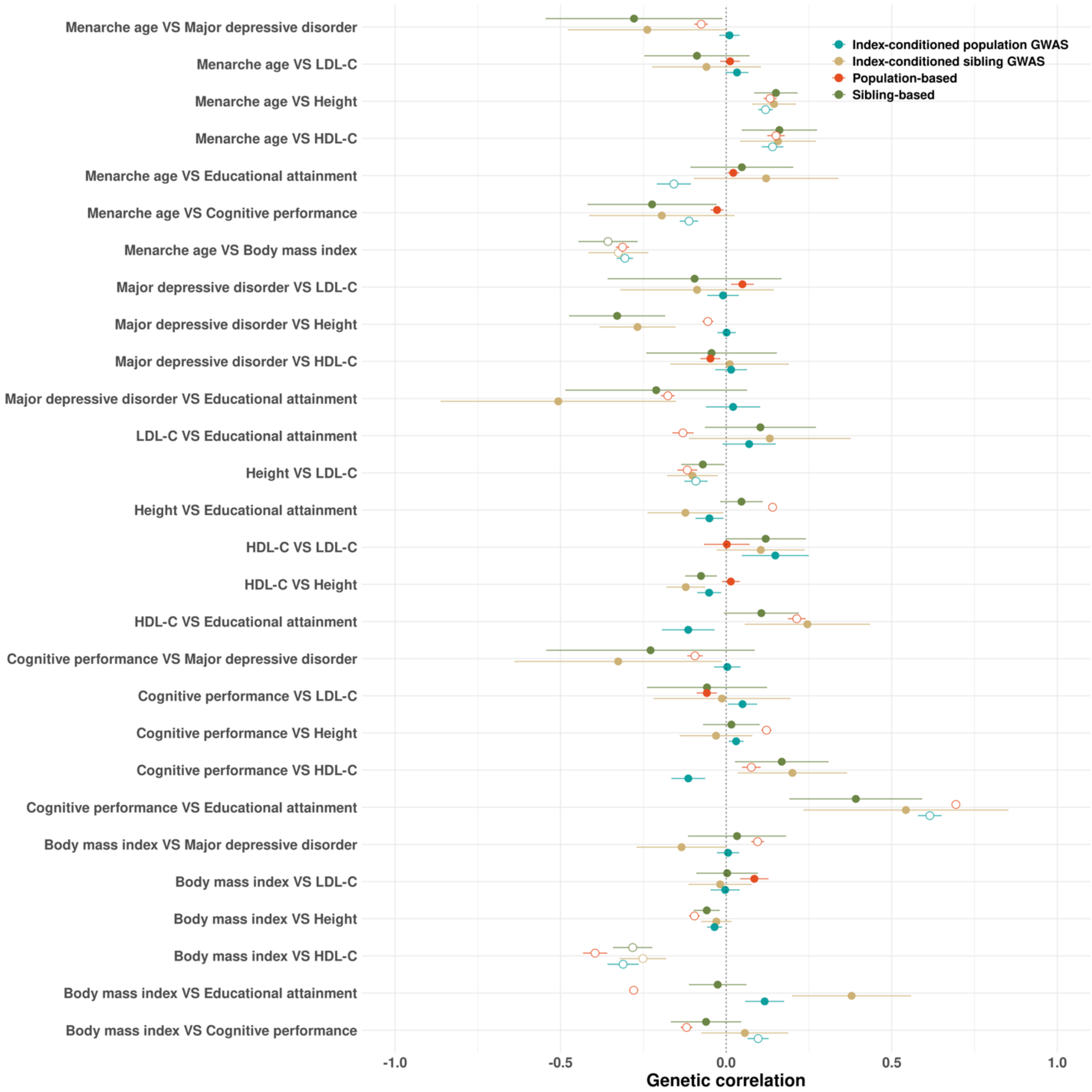


**Supplementary Figure 18. Genetic correlation estimates based on population GWAS, conditional analysis, and within-sibling GWAS.** Hollow circles indicate significant correlations at FDR < 0.05. Intervals indicate standard errors.

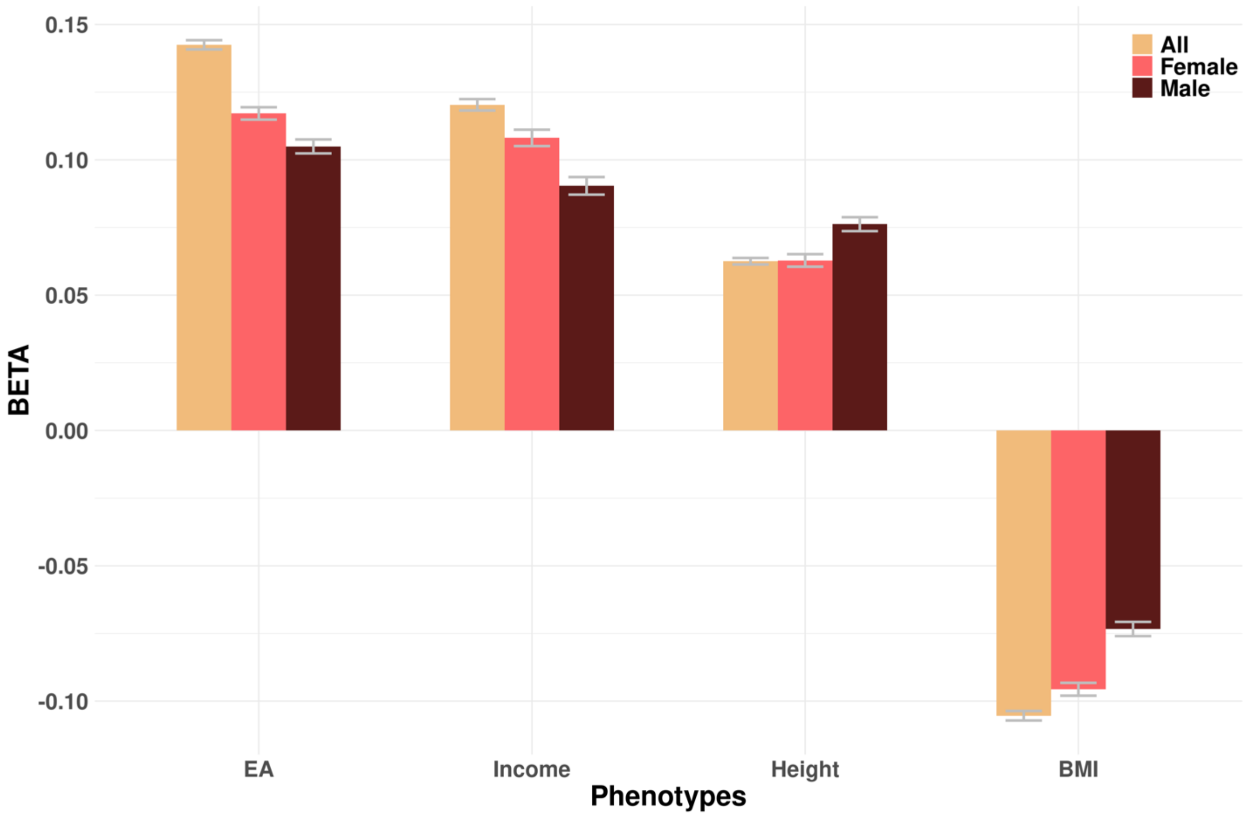


**Supplementary Figure 19. Association between the index PGS and four phenotypes in UKB.** Each bar shows the index PGS-phenotype association after controlling for sex, age, and genetic principal components. Error bars are standard errors.

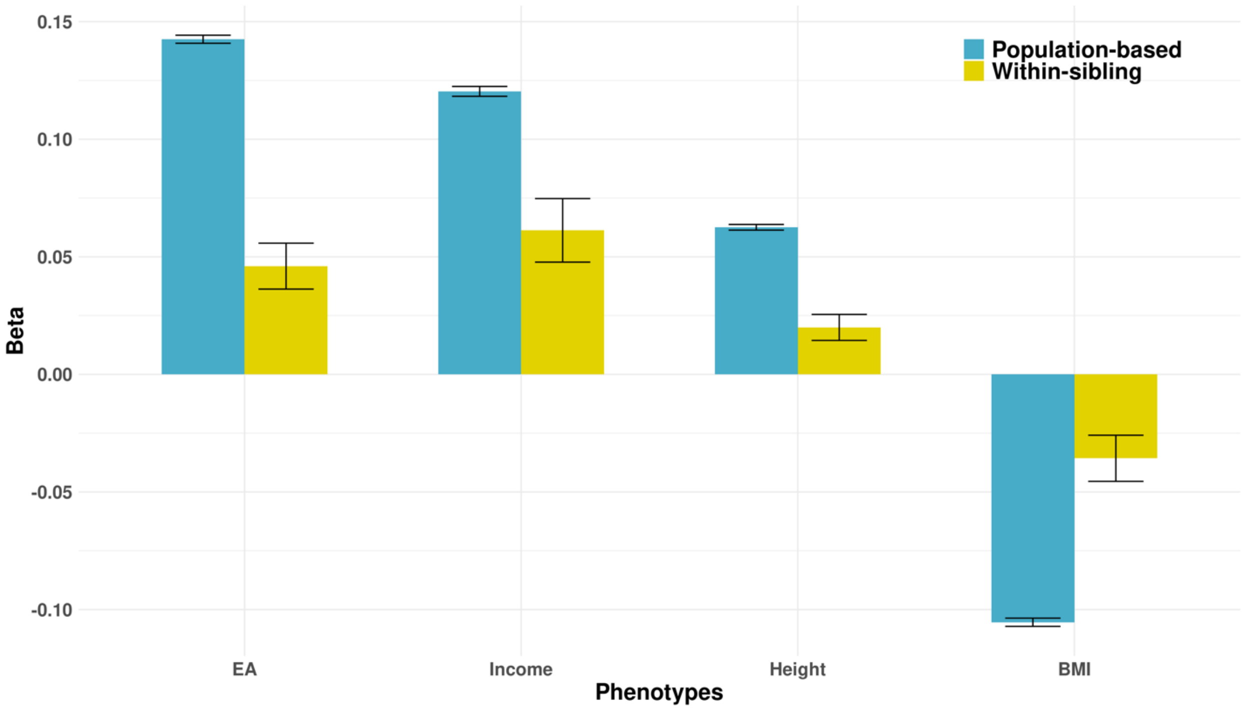


**Supplementary Figure 20. PGS regression coefficients estimated from population-based and within-sibling analyses.** Error bars are standard errors.

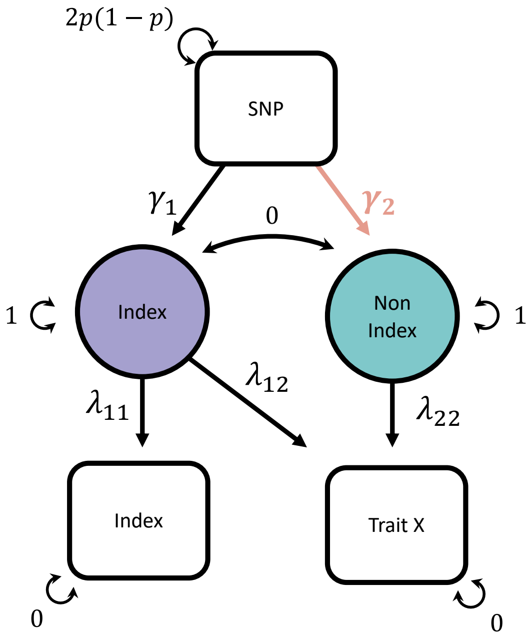


**Supplementary Figure 21. Schematic diagram for using GWAS-by-subtraction to reduce assortative mating-driven biases in genetic associations.** The main parameter of interest, i.e., genetic effect on the non-index-driven component, $\gamma_2$, for each trait X, is highlighted in light pink.
