## Supplementary Note for "Genetic basis of partner choice"

### General model setting

We assume that the multi-dimensional individual preferences in partner choice can be summarized by a one-dimensional index which is a function of many characteristics[1] (e.g., education, income, physical attractiveness).

$$I^m = f(X_1, X_2, \dots, X_K)$$
$$I^f = g(Y_1, Y_2, \dots, Y_L)$$

This model assumes that an individual's characteristics matter for the partner's preference only through the index. Here, notations are consistent with the main text except that $f$ and $g$ can be nonlinear functions. In the economics literature, it is often important to distinguish observable and unobservable characteristics. This is not crucial in our study. The reason is that we think of each of these characteristics as a function of genetic and environmental variables. Genetic variables are always observable even when a subset of characteristics is not. The relationship between a man's (woman's) characteristics $X$ $(Y)$, SNP genotypes $G$ $(S)$, and non-genetic factors $\varepsilon$ $(\delta)$ can be generalized as follows

$$X_k = \pi_k(G, \varepsilon_k), k = 1, \dots, K$$
$$Y_l = \rho_l(S, \delta_l), l = 1, \dots, L$$

In the main text, $\pi$ and $\rho$ are linear, a standard assumption in GWAS studies.

Our main results depend on two key assumptions:

- ***Separability*** holds if genetic variants only matter for partner choice through genetic components $\tilde{I}^m = \theta(G)$ and $\tilde{I}^f = \tau(S)$:
$$I^m = f(X) = \tilde{f}(\tilde{I}^m, \varepsilon) = \tilde{f}(\theta(G), \varepsilon)$$
$$I^f = g(Y) = \tilde{g}(\tilde{I}^f, \delta) = \tilde{g}(\tau(S), \delta).$$
- ***Conditional Independence*** holds if the distribution of $\varepsilon$ $(\delta)$ is independent of $G$ $(S)$ conditional on $\tilde{I}^m(\tilde{I}^f)$.

Our main text has provided details under the linear model setting. Here, linearity is a sufficient but not necessary condition for these assumptions to hold.

Under the separability and conditional independence assumptions, a stable matching in the marriage market exists such that the distribution of female characteristics $(Y_1, \dots . Y_L)$ conditional on the characteristics of her spouse $(X_1, \dots . X)$ depends only on the partner choice index of his spouse, $I^m$, and vice versa (see Chiappori et al. 2012 for a proof)[1]. Given the conditional independence assumption, it also follows that the conditional distribution of female characteristics $(Y_1, \dots . Y_L)$ depends on the male SNP genotypes $G$ only through the genetic component of $I^m$. Crucially, this property extends to all moments of the conditional distribution of $(Y_1, \dots . Y_L)$, and in particular we have:

$$E(Y_l \mid G) = E(Y_l \mid \tilde{I}^m) = \varphi_l(\tilde{I}^m)$$

That is, the conditional expectation of the l-th characteristic of the wife given the husband's genetics $G$ is a function of the husband's index $I^m$. A similar conclusion holds true for the wife's index $\tilde{I}^f$. Therefore, although we cannot generally identify $I^m(I^f)$, its genetic components $\tilde{I}^m(\tilde{I}^f)$ can be identified (up to some transformation) using some couples' observable characteristics. Further, the relative contribution of genetic variant $G_j$ on the genetic component $\tilde{I}^m$ (i.e., $\partial\tilde{I}^m/\partial G_j$) can be empirically studied.

$$\frac{\partial\mathbb{E}(Y_l|G)}{\partial G_j} = \frac{\partial\varphi_l}{\partial\tilde{I}^m} \times \frac{\partial\tilde{I}^m}{\partial G_j}$$

Note that $\partial\varphi_l/\partial\tilde{I}^m$ does not depend on $j$. If we impose that $\partial\varphi_l/\partial\tilde{I}^m$ is a constant for different values of $\tilde{I}^m$, $\partial\tilde{I}^m/\partial G_j$ can be obtained from empirically observable $\partial\mathbb{E}(Y_l|G)/\partial G_j$ up to a constant.

**Some discussion on the separability assumption**

We have demonstrated in the main text that both separability and conditional independence assumptions would hold under a simple linear model setting. Here, we provide additional discussions and insights on the separability assumption, by illustrating two examples, one where the separability assumption holds under different model assumptions than in the main text, and another where it does not.

*Example 1.* Suppose the index $I^m(I^f)$ has with the following structure:

$$I^m = \prod_j^M G_j \prod_k^K \varepsilon_k$$

$$I^f = \prod_j^M S_j \prod_l^L \delta_l$$

Then, the SNP genotypes $G(S)$ matter through a single factor $\tilde{I}^m$ ($\tilde{I}^f$) and the non-genetic component $\varepsilon(\delta)$ matters separately. The genetic components $\tilde{I}^m$ and $\tilde{I}^f$ correspond to:

$$\tilde{I}^m = \prod_j^M G_j$$

$$\tilde{I}^f = \prod_j^M S_j$$

Therefore, under this setting, the separability assumption would hold. However, this structure is not consistent with the initial idea that both genetic and non-genetic components matter through a set of characteristics $X(Y)$. That is, $f$ and $g$ are undefined. Although our main theoretical results will hold in this setting, we would not be able to obtain any empirical findings, as conducting a GWAS on $I^m$ ($I^f$) directly is unfeasible, given that these indices are latent.

*Example 2.* Another example is an index $I^m(I^f)$ with the following structure:

$$I^m = \prod_k^K X_k = \prod_k^K \pi_k(G, \varepsilon_k)$$

$$I^f = \prod_l^L Y_l = \prod_l^L \rho_l(S, \delta_l)$$

In this case, the separability assumption would not be respected even if each characteristic $X_k(Y_l)$ linearly depends on genes and environments (i.e., $\pi_k$ and $\rho_l$ are linear functions). The key problem here is that the same genes matter through different characteristics $X_k(Y_l)$ which are also partly shaped by non-genetic components.

However, although we assumed in the main text that each characteristic is a linear combination of genetic and non-genetic components, nothing prevents us from specifying the characteristics $X_k(Y_l)$ in a flexible manner

$$X_k = \sum_{j=1}^{M} G_j \beta_{jk} + \varepsilon_k, \qquad k = 1, \dots, K$$

$$Y_l = \sum_{j=1}^{M} S_j \gamma_{jl} + \delta_l, \qquad l = 1, \dots, L$$

For example, $X_k$ could be either income, log-transformed income, or even income rank in the distribution. Quadratic terms of BMI could be modeled to represent a U-shaped relationship between BMI and the index; alternatively, one could use dummy variables to flag BMI categories (e.g., underweight, overweight, obesity). EA can also be decomposed into categorical dummy variables (e.g., college or high school attainment), each represented by a different element of $X$.

**Identifying the shared genetic component across multiple spousal traits**

Our main results have demonstrated that genetic associations with the index $I^m(I^f)$ under the linear model setting, can be proportionally identified through GWAS on spousal traits. Importantly, the choice of spousal trait $Y_l$ does not have a major impact on the results. However, in practice, we may consider using multiple spousal traits to improve the analysis. The reason of this is two-fold. First, we may come up with more precise estimates of genetic associations by exploiting the additional variation across different (spousal) traits. This is analogous to performing a meta-analysis to combine multiple sets of GWAS associations[2]. Second, and more importantly, this allows us to obtain more robust association results even when the index-based matching model is partially violated.

While the single-index model provides very nice theoretical properties, it is an over-simplification of the complex nature of marriage market. The single index model presumes that all males (females) in the population choose their spouse by weighting various characteristics of potential spouses in a similar way. That is, in the single index model, positive correlations of traits among couples are induced because males that are considered attractive by other females are more likely to couple with females that are considered attractive by other males, and vice versa. For example, if a college graduate is perceived to be more attractive by both those with and without a college degree, EA behaves in accordance to the single index model. In practice, however, individual traits may undergo additional sorting that is not explained by a single index alone (**Supplementary Figure 1**). For example, college-educated individuals are more likely to meet their partner in environments where college graduates are overrepresented, which will lead to a spousal correlation in EA even if higher EA does not make someone more attractive. Another example is that if a college graduate would prefer to marry another college graduate, whereas those without a college degree would prefer to marry someone without a college degree, then EA does not behave according to the single index model. We call such instances trait-specific sorting. Note that, both under index-driven sorting and trait-specific sorting, a positive correlation of EA is observed among couples. Another example from a recent empirical investigation is that income behaves according to the single index model, but smoking acts differently, with non-smokers preferring non-smokers and smokers preferring smokers[3]. When trait-specific sorting mechanisms exist, the relationship between husband's genetics and wife's characteristic, for example EA, is no longer explained by husband's index alone. Instead, husband's EA needs to be considered as well. Here, without loss of generality, suppose that $Y_1(X_1)$ denotes wife's (husband's) EA. Then we have:

$$E(Y_1 \mid G) = E(Y_1 \mid \tilde{I}^m, X_1) = \tilde{\varphi}_1(\tilde{I}^m, X_1)$$

Therefore, SNP effect estimates obtained in spousal EA GWAS are mixtures of genetic associations with the index $\tilde{I}^m$ (i.e., $\partial \tilde{I}^m / \partial G_j$) and genetic associations with own EA (i.e., $\partial X_1 / \partial G_j$).

$$\frac{\partial \mathbb{E}(Y_1|G)}{\partial G_j} = \frac{\partial \tilde{\varphi}_1}{\partial \tilde{I}^m} \times \frac{\partial \tilde{I}^m}{\partial G_j} + \frac{\partial \tilde{\varphi}_1}{\partial X_1} \times \frac{\partial X_1}{\partial G_j}$$

Under this setting, even under linear assumptions, GWAS of a single spousal trait alone is no longer proportional to GWAS of the partner choice index. However, different spousal trait GWAS will all share the same linear component representing index genetic associations. When the own traits' true genetic associations do not share a pleiotropic component (i.e., $\partial X_1 / \partial G_j$ do not have shared genetics across traits), factor analysis aiming to identify the shared linear component underlying various spousal trait GWAS will recover the index GWAS up to a linear transformation. This can be achieved by including a set of traits with distinctive genetic basis into the analysis (e.g., height and cognition). On the other hand, if the analysis were based on a set of (spousal) psychiatric outcomes that share psychopathology factor, index GWAS will not be correctly identified.

This is the motivation behind GenomicSEM factor GWAS in our study (see **Figure 3A** in the main text). We use GenomicSEM to identify the shared genetic component across multiple spousal traits and use it to represent the GWAS of partner choice index. The trait-specific genetic components that cannot be explained by the index will be absorbed into the residual variance of spousal traits in this model.

Another potential solution to this problem is to adjust for own trait as a covariate when performing GWAS of a spousal trait. This will allow us to remove the biases from own trait genetic associations from the GWAS estimates.

$$\frac{\partial \mathbb{E}(Y_1 \mid G, X_1)}{\partial G_j} = \frac{\partial \tilde{\varphi}_1}{\partial \tilde{I}^m} \times \frac{\partial \tilde{I}^m}{\partial G_j}$$

However, we expect this approach to substantially reduce statistical power in genetic association tests.
